# Molecular determinants of brain-resident CD8^+^ T cell formation and function

**DOI:** 10.1101/2025.08.26.672425

**Authors:** Tarek Elmzzahi, Chun-Hsi Su, Mehrnoush Hadaddzadeh Shakiba, Doaa Hamada, Darya Malko, Maren Köhne, Aleksej Frolov, Teisha Mason, Rebekka Scholz, Yuanfang Li, Collins Osei-Sarpong, Leonie Heyden, Jonas Schulte-Schrepping, Lorenzo Bonaguro, Kristian Händler, Vassiliki A. Boussiotis, Annett Halle, Elena De Domenico, Daniel Gray, Martin Fuhrmann, Zeinab Abdullah, Axel Kallies, Kevin Man, Marc D. Beyer

**Author notes:** These authors jointly supervised this work (Axel Kallies, Kevin Man, Marc D. Beyer).

## Abstract

Tissue-resident memory T (Trm) cells are strategically located to provide frontline protection upon antigen re-encounter while possessing tissue-specific transcriptional programs. Whether brain Trm cells similarly adapt to their tissue environment, and to what extent their molecular signature is altered in neuropathology, remains unclear. Here we profile brain Trm cells under homeostasis and in the contexts of aging, beta-amyloidosis, and systemic viral infection. From these studies, a tissue-specific CD8^+^ T cell landscape emerged, defined by the expression of the transcription factor TCF-1 and the inhibitory receptor PD-1. TCF-1 was critical for the formation and phenotypic maturation of brain CD8^+^ Trm cells, while PD-1 signaling was necessary for robust effector function and antigen-specific recall response. In addition, the cytokine transforming growth factor (TGF)-β was required for the differentiation of brain CD8^+^ Trm cells and restricted their transition into effector-like cells upon antigenic rechallenge. These findings highlight common as well as tissue-specific features of brain CD8^+^ Trm cells and provide insights into the molecular mechanisms governing their formation and function.

## Introduction

Compared to their naive counterparts, memory T cells exhibit a more rapid and profound effector response upon secondary antigen encounter.^1,2^ Memory CD8^+^ T cells are broadly categorized into circulating and non-recirculating, tissue-resident memory T (Trm) cells.^3,4^ Circulating memory T cells are classified into different subsets, namely central and effector memory T cells as well as CX3CR1^+^ peripheral memory T cells, based on their surface phenotype, longevity, migration pattern, and capacity for recall.^5–8^ Trm cells are strategically located at sites of previous insult to confer frontline protection and are poised for a rapid recall response upon antigen re-encounter.^9^ Trm cells also possess a superior cytolytic capacity compared to their circulating memory counterparts.^3,9^ An additional facet of Trm cell responses is the innate-like induction of a pro-inflammatory “alarm state” in the responding tissue,^10–12^ which facilitates the recruitment of circulating innate and adaptive immune cells.^11^ Overall, Trm cells mediate host protection through T cell intrinsic and extrinsic mechanisms.^13^

CD8^+^ Trm cell differentiation is initiated during the early stages of the immune response, with memory-precursor T cells seeding non-lymphoid tissues by day 4.5 to day 7 post-infection.^14,15^ Mature CD8^+^ Trm cells are transcriptionally and phenotypically distinct compared to circulating memory T cells.^16,17^ Numerous studies have characterized Trm cells using surface markers such as CD69, the integrin α chains CD103 and CD49a, the chemokine receptor CXCR6, as well as transcription factors including Runx3, Blimp-1, and Hobit.^9,15,18,19^ Importantly, while Trm cells across different tissues share key transcriptional features,^19^ they do not exhibit a uniform molecular profile.^20,21^ Indeed, high-dimensional analysis of Trm cells across tissues revealed that the tissue of residence exerts a critical role in shaping the molecular hallmarks of Trm cells.^20,21^ This influence is attributed to the presence or absence of local antigen as well as microenvironmental cues that drive distinct Trm cell phenotypes.^9^ One important cytokine underpinning this local tissue adaptation is transforming growth factor (TGF)-β. TGF-β is crucial for the development and maintenance of Trm cells in skin,^17,22^ small intestine,^23,24^ and salivary gland,^25^ and regulates the expression of CD103 and CD49a.^26^ To what extent the tissue microenvironment is a strong determinant of brain CD8^+^ Trm cell differentiation, and whether TGF-β shapes brain CD8^+^ Trm cell formation, remain unknown.

Early studies investigating brain CD8^+^ Trm cells have mostly employed direct brain infection via intracranial or intranasal delivery of pathogens.^27,28^ Brain CD8^+^ Trm cells persist without replenishment by circulating memory T cells or ongoing antigen encounter, and autonomously mediate pathogen clearance upon antigenic rechallenge.^27,29^ The majority of such CD8^+^ Trm cells express CD103, which is necessary to promote T cell retention in the brain.^27,29^ Notably, intraperitoneal infection with lymphocytic choriomeningitis virus (LCMV)-Armstrong does not manifest in brain infection^30^ yet results in a brain CD8^+^ T cell population that exhibits hallmarks of tissue residency, including expression of CD69 and CD49a and resistance to systemic depletion using anti-CD8α antibody.^31^ However, <10% of brain CD8^+^ Trm cells express CD103 in this context,^31^ highlighting the requirement for local antigen recognition during priming to induce CD103 expression in brain CD8^+^ Trm cells.^28^ Importantly, such “peripherally induced” brain CD8^+^ Trm cells were sufficient to confer protection against local antigenic rechallenge independent of the contribution of circulating memory T cells.^31,32^

Recent studies have employed single-cell RNA-sequencing (scRNA-seq) to decipher the molecular landscape of brain T cells in the contexts of ageing,^33^ beta-amyloidosis,^34^ tauopathy,^35^ and autoimmunity.^36^ However, it remains unclear whether the transcriptional makeup of brain T cells is underpinned by the (patho)physiological context or brain tissue residency *per se*. In addition, the majority of the aforementioned studies employed either constitutive gene deletion (*Cd8* or *Rag2* KO mice), or depletion of circulating CD8^+^ T cells to assess the role of CD8^+^ T cells. Accordingly, the impact of the tissue microenvironment on the differentiation of brain CD8^+^ Trm cells, and the signaling pathways and transcription factors regulating their formation and function, warrant further investigation.

In this study, we identify molecular determinants of the differentiation and function of brain Trm cells, with a particular focus on CD8^+^ T cells. Our data reveal that brain CD8^+^ Trm cells acquire a primarily tissue-dependent rather than context-dependent phenotype, with similar molecular hallmarks observed across steady state, ageing, beta-amyloidosis, and post-acute but not chronic viral infection. Molecularly, we observed that TCF-1, PD-1, and TGF-β regulate the differentiation and functional properties of brain CD8^+^ T cells. Overall, our data reveal common as well as tissue- and context-specific features of brain CD8^+^ Trm cells and provide novel insights into the molecular mechanisms that guide their formation and function.

## Results

### TCF-1 and PD-1 define developmentally and functionally distinct brain CD8^+^ Trm cell subsets

T cells populate the brain at steady state,^37–40^ but the transcriptional and phenotypic diversity among such cells remains poorly understood. To unravel the complexity of the brain T cell landscape during homeostasis, we sorted CD3^+^ T cells from the brains of 5-month-old naive mice – after exclusion of circulating T cells via anti-CD45 intravascular labelling^41^ – and examined their transcriptome using scRNA-seq. Unbiased clustering revealed multiple subsets of αβ T cells as well as unconventional T cells (**Figures 1A** and **1B**). Four main clusters of CD8^+^ T cells were identified. One CD8^+^ cluster, defined by *Pdcd1* (encoding PD-1), exhibited high expression of transcripts encoding granzyme K (*Gzmk*), chemokines (*Ccl5*, *Ccl4*) and transcription factors associated with T cell receptor (TCR) signaling (*Tox*, *Nr4a2*). Another CD8^+^ cluster was marked by high expression of *Ly6c2* (encodes Ly6C), as well as *Lgals3* and the transcription factor *Hopx*. A third cluster showed a pronounced signature of type I interferon (IFN)-stimulated genes (*Isg*) and IFN-induced protein with tetratricopeptide repeats (IFIT) genes. In addition, we observed a subset of T cells that expressed transcripts encoding surface molecules and transcription factors associated with central memory (Tcm) or stem-like T cells, including *Sell* (encoding CD62L), and *Tcf7* (encoding TCF-1). Furthermore, small clusters of CD4^+^ T cells, γδ T cells, and NKT cells were identified, in agreement with previous reports.^38,40^

**Figure 1.**
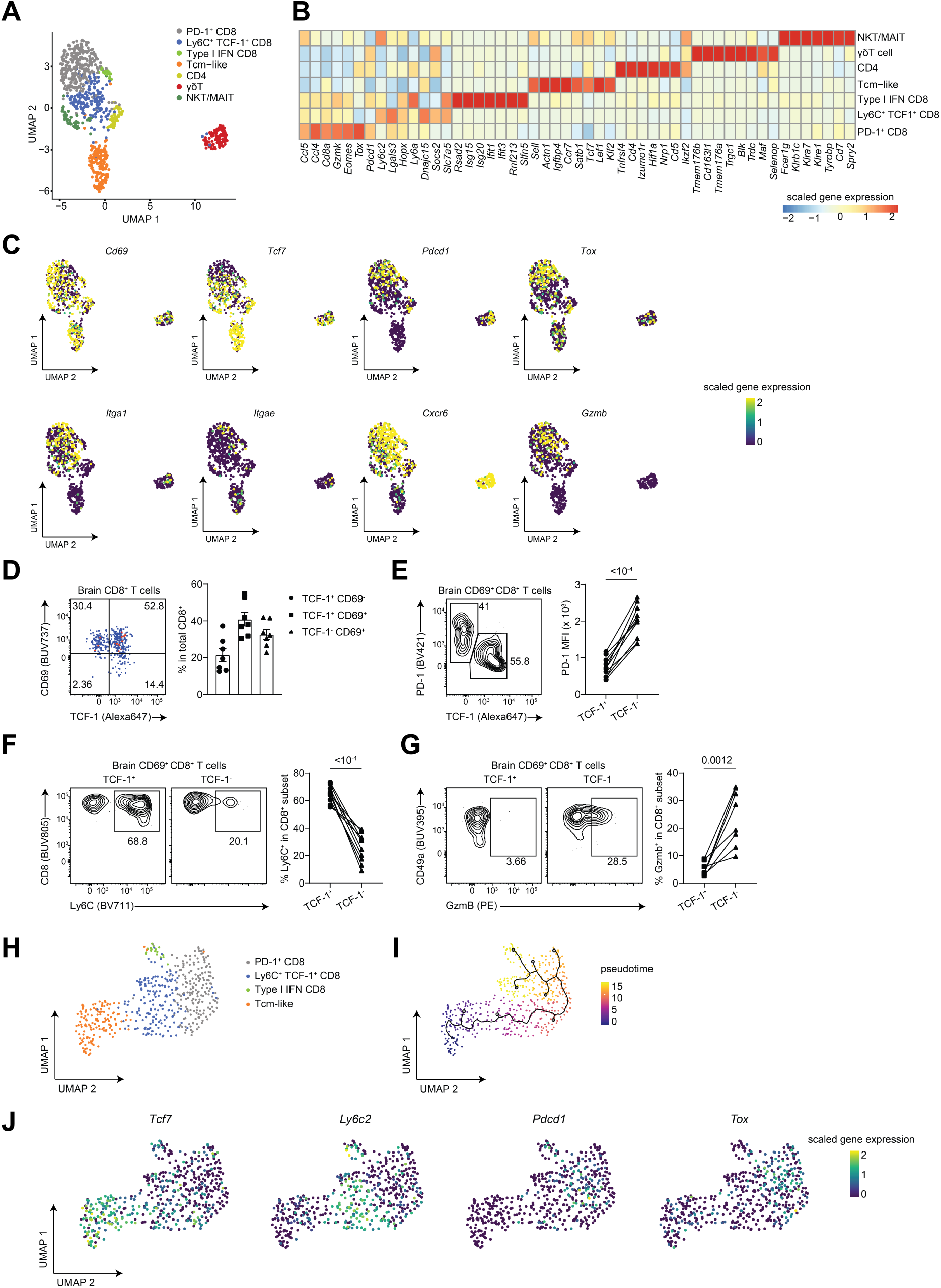
Transcriptional and phenotypic heterogeneity of the brain T cell compartment at steady state. **A**, Uniform Manifold Approximation and Projection (UMAP) plot of scRNA-seq analysis of extravascular CD3^+^ T cells (n = 798 cells) in the brain of 5 month-old mice; n = 5 mice. Cells were processed using the BD Rhapsody 3’ protocol. **B**, Heatmap of the top differentially expressed genes defining each T cells cluster. **C**, FeaturePlots of a select set of genes overlaid on the UMAP described in **A**. **D**, flow cytometric analysis and quantification of frequencies of brain CD8^+^ T cells based on CD69 and TCF-1 coexpression; n = 7 mice. **E**, PD-1 and TCF-1 co-expression in brain CD69^+^ CD8^+^ T cells (left) and PD-1 MFI in the respective populations (right); n = 7 mice. **F**, Ly6C expression in TCF-1^+^ and TCF-1^-^ subsets of brain CD69^+^ CD8^+^ T cells; n = 10 mice. **G,** proportion of GzmB^+^ in TCF-1^+^ and TCF-1^-^ subsets of brain CD69^+^ CD8^+^ T cells; n = 8 mice. **H**, CD8^+^ T cell clusters from the scRNA-seq dataset described in a were subsetted and re-analyzed; 527 cells; n = 6 mice. **I**, monocle3 trajectory inferred across the CD8^+^ T cell clusters; blue represents least differentiated and yellow represents terminally differentiated. **J**, FeaturePlots of a number of genes marking the CD8^+^ T cell subsets. Data are pooled (D-G) from two independent experiments. Statistical analyses were performed using paired (E,F) or unpaired (G) two-tailed Student’s *t* test. Error bars represent standard error of the mean (SEM). GzmB, granzyme B; MFI, median fluorescence intensity; Tcm, central memory T cells.

As expected, *Cd69* was expressed by the majority of brain Trm cells (**Figure 1C**). Similarly, transcripts encoding the chemokine receptor CXCR6 and cytotoxic molecule granzyme B (GzmB) were produced by a large fraction of brain CD8^+^ T cells (**Figure 1C**). *Itgae* (encoding CD103) was co-expressed with *Cd69* by only ∼10% of brain CD8^+^ Trm cells, supporting the notion that the CD69-expressing CD8^+^ T cells developed independent of CD103 expression (**Figure 1C** and **Figure S1A**). In contrast, *Itga1* (encoding the TGF-β-dependent CD49a) and *Cd69* were co-produced by >60% of brain CD8^+^ T cells (**Figure 1C** and **Figure S1A**).

The transcription factor TCF-1 regulates the development of naive and Tcm cells;^42,43^ however, its function in Trm cells is poorly understood. Interestingly, a discernible subset of brain CD8^+^ T cells displayed co-expression of CD69 and TCF-1 or PD-1, respectively, at both the transcriptional and protein levels (**Figures 1C-1E**). PD-1^-^ TCF-1^+^ CD69^+^ CD8^+^ T cells (TCF-1^+^ CD8^+^ Trm cells) co-expressed Ly6C and mostly corresponded to the *Ly6c2*^+^ CD8^+^ cluster, whereas PD-1^+^ TCF-1^-^ CD69^+^ CD8^+^ T cells (TCF-1^-^ CD8^+^ Trm cells) largely represented *Pdcd1*^+^ cells in the scRNA-seq data (**Figure 1F** and **Figure S1B**). Brain TCF-1^+^ Trm cells exhibited greater proliferation but produced less GzmB and IFNγ (**Figure 1G** and **Figures S1C** and **S1D**), consistent with the reported inverse correlation between TCF-1 and GzmB expression.^42,44^ Thus, TCF-1 and PD-1 mark two phenotypically distinct subsets of CD69^+^ CD8^+^ Trm cells in the brain.

To investigate the developmental hierarchy among brain CD8^+^ T cell subsets, we subsetted the CD8^+^ T cell clusters that are part of the scRNA-seq dataset described above (**Figure 1H**) and performed pseudotime analysis using monocle3 (ref. 45). This analysis revealed a differentiation trajectory starting from Tcm-like cells and progressing through *Pdcd1*^+^ cells towards the type I IFN cluster (**Figure 1I**). This trajectory was associated with progressive *Tcf7* downregulation and a reciprocal upregulation of *Pdcd1* and *Tox* (**Figure 1J**).

Collectively, these findings indicate that brain CD8^+^ T cells are heterogeneous and bear transcriptional and phenotypic hallmarks of Trm cells in peripheral non-lymphoid tissues. Furthermore, these data suggest that brain CD8^+^ Trm cells undergo a stepwise differentiation from a TCF-1^+^ state to a TCF-1^-^ PD-1^+^ state with progressive acquisition of effector function.

### Ageing-induced quantitative but not qualitative alterations in brain CD8^+^ Trm cells

Ageing is associated with profound systemic changes in the T cell compartment in lymphoid and non-lymphoid tissues, including upregulation of PD-1 (ref. 46,47). We therefore asked how ageing modulates the brain CD8^+^ T cell compartment. Old mice (>21 months of age) exhibited substantially larger numbers of brain CD8^+^ T cells compared to their young counterparts (**Figure 2A**). Numerically, both TCF-1^+^ and TCF-1^-^ brain CD8^+^ Trm cells were increased in old mice, with no difference in the frequencies of these subsets (**Figure 2B**). Furthermore, there were no differences in the frequencies of PD-1^+^ cells among brain CD8^+^ Trm cells in young and old mice (**Figure 2C**).

**Figure 2.**
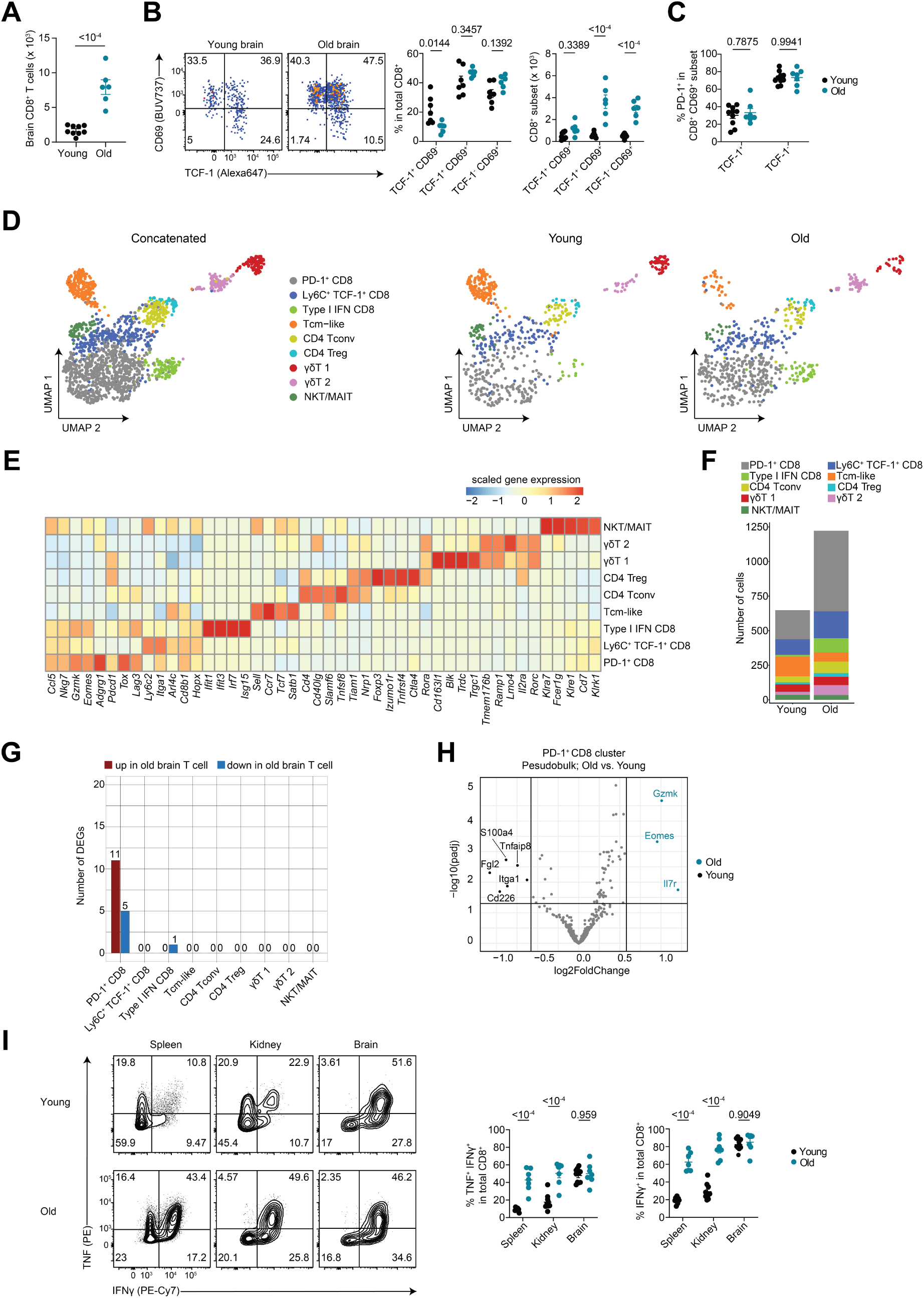
Ageing-induced alteration of the brain CD8^+^ T cell compartment is primarily quantitative rather than qualitative. **A**, quantification of brain CD8^+^ T cells in young (5-7 months) and old (21-24 months)-old naive, sex-matched mice; n = 6-9 mice per group. **B**, frequencies and numbers of TCF-1^+^ CD69^-^, TCF-1^+^ CD69^+^, and TCF-1^-^ CD69^+^ subsets of brain CD8^+^ T cells in young and old mice; n = 6-9 mice per group. **C**, percentages of PD-1^+^ among TCF-1^+^ and TCF-1^-^ subsets of brain CD69^+^ CD8^+^ T cells; n = 7-10 mice per group. **D**, Uniform Manifold Approximation and Projection (UMAP) plot of scRNA-seq analysis of extravascular CD3^+^ T cells (n = 1870 cells) in the brain of 5 month-and 21 month-old naive mice; n = 5 mice per group. Cells were processed using the BD Rhapsody 3’ protocol. **E**, heatmap showing the top differentially expressed (DE) genes defining each T cell cluster. **F**, barplot showing the number of cells per T cell cluster in young and old mice. **G**, barplot showing the number of DE genes in a pairwise comparison between young and old mice per T cell cluster. **H**, volcano plot depicting the DE genes between cells corresponding to PD-1^+^ CD8 T cell cluster in young vs. old mouse brain; DE genes were computed using a pseudobulk approach. **I**, frequencies of TNF^+^ IFNγ^+^ and total IFNγ^+^ CD8^+^ T cells in the spleen, kidney, and brain of young (6-7 month) and old (24 month) mice following *ex vivo* stimulation using PMA/ionomycin; n = 7-10 mice per group. Data are pooled (A-C,I) from at least two independent experiments. Statistical analyses were performed using unpaired two-tailed Student’s *t* test (A) or two-way ANOVA with Šídák’s multiple comparisons test (B,C,I).

To further explore potential age-induced alterations of the brain T cell landscape, we sorted CD3^+^ cells from young and old mouse brains and performed scRNA-seq. Interestingly, the global composition of the T cell compartment in aged mice was similar to that of young mice: we observed four main subsets of CD8^+^ T cells, which were transcriptionally similar to their counterparts in young mice (**Figures 2D** and **2E**). Ageing, however, was associated with altered frequencies of these clusters, with notable increases in numbers of PD-1^+^ cells and type I IFN CD8^+^ T cells (**Figure 2F**). To address whether ageing induces a qualitative alteration in the gene expression program of brain T cells, we performed pairwise differential gene expression (DE) analysis for each brain T cell cluster across young and old mice. Remarkably, the majority of brain T cell clusters showed no differentially expressed genes between young and old mice (**Figure 2G**), except for PD-1^+^ CD8 T cells, which exhibited a relatively small total number of 16 DE genes (**Figure 2G**). Specifically, signature genes associated with the PD-1^+^ CD8 T cell cluster were upregulated in PD-1^+^ CD8 Trm cells from aged compared to young mice, suggesting age-induced maturation of PD-1^+^ CD8^+^ Trm cells (**Figure 2H**).

Notably, brain CD8^+^ Trm cells from young and old mice displayed similar capacities to express TNF and IFNγ following *ex vivo* stimulation (**Figure 2I**). This was in contrast to splenic and renal CD8^+^ T cells, which, consistent with previous studies,^46^ exhibited an increased frequency of TNF^+^ IFNγ^+^ and total IFNγ^+^ CD8^+^ T cells in old compared to young mice (**Figure 2I**).

Collectively, these data demonstrate that the mouse brain supports the accumulation of CD8^+^ Trm cells in an age-dependent manner, while transcriptional, phenotypic and functional properties of such cells are imprinted in a tissue-specific rather than age-dependent fashion.

### Tissue-specific rather than disease-specific imprinting of brain CD8^+^ Trm cells

Alzheimer’s disease (AD) is associated with profound alterations of the central and peripheral immune compartments.^48^ We evaluated whether beta-amyloidosis, a hallmark of AD neuropathology, affects the composition and transcriptional profile of brain Trm cells. To this end, we made use of the APP/PS1dE9 mouse model of beta-amyloidosis.^49^ As expected, a subset of microglia in APP/PS1dE9 mice exhibited a “disease-associated microglia” phenotype,^50,51^ as defined by increased CD11c expression (**Figure S2A**). Consistent with a previous report,^52^ the number of brain CD8^+^ T cells showed a gradual, age-dependent increase in APP/PS1dE9 mice, with a distinct accumulation observed in 15-month-old APP/PS1dE9 mice (**Figure 3A**). Intriguingly, this increase was selectively attributed to an increase in TCF-1^+^ CD69^+^ CD8^+^ Trm cells (**Figures 3B** and **3C**). We next focused on the 10-month and 15-month time-points and used scRNA-seq to evaluate whether the increase in CD8^+^ T cell numbers was accompanied by transcriptional alterations (**Figures 3D-F** and **Figures S2B-E**). Consistent with our observations in wildtype (WT) mice, we identified 4 clusters of CD8^+^ T cells, defined largely by the expression of *Pdcd1*, *Ly6c2*, type I IFN response genes, and a Tcm phenotype (**Figures 3D** and **3E**). Pairwise analysis of DE genes for each cluster revealed almost no difference in the transcriptional makeup of the different T cell subsets in APP/PS1dE9 compared to age-matched control mice (**Figure 3G** and **Figure S2E**), supporting the notion that brain CD8^+^ T cells acquire a tissue-specific transcriptional signature. Similarly, there were no differences in the frequencies of TNF^+^ IFNγ^+^ or total IFNγ^+^ brain CD8^+^ T cells between APP/PS1dE9 and control mice (**Figure 3H**), although there was a moderate increase in the amount of IFNγ produced by CD8^+^ T cells in APP/PS1dE9 mice (**Figure 3H**). To validate our findings in an additional model of accelerated beta-amyloidosis, we made use of 5xFAD mice.^53^ Similar to our observation in APP/PS1dE9 mice, the total number of CD8^+^ T cells was increased in the brains of 9-month-old 5xFAD mice compared to controls (**Figure S2F**). This increase was also selectively attributed to the TCF-1^+^ CD69^+^ subset of brain CD8^+^ Trm cells (**Figure S2G**). In addition, brain CD8^+^ T cells in 5xFAD mice produced more IFNγ upon *ex vivo* stimulation (**Figure S2H**).

**Figure 3.**
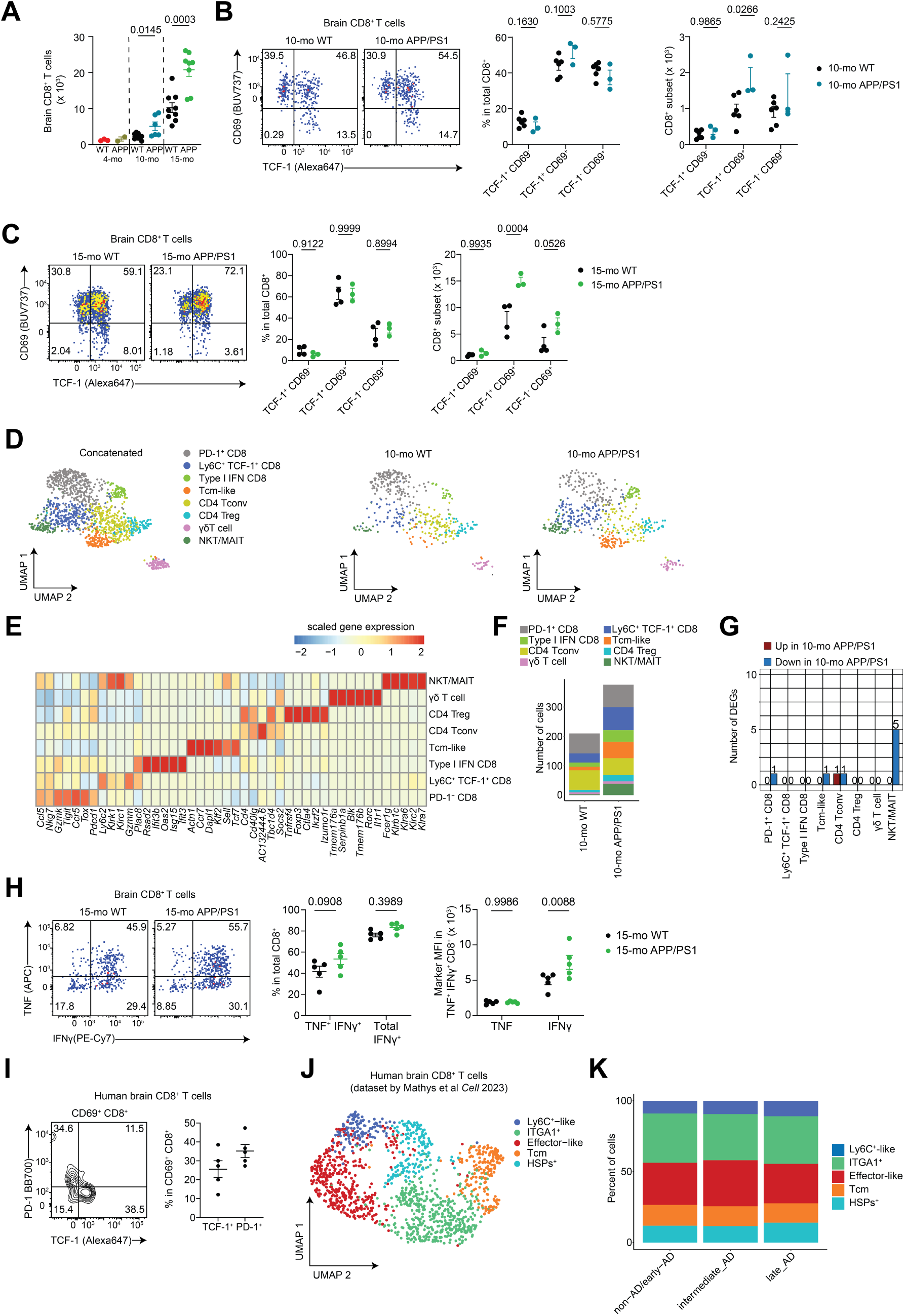
Tissue-specific rather than disease-specific imprinting of brain CD8^+^ Trm cells in neurodegeneration. APP/PS1dE9 transgenic mice and sex-and age-matched mice were used for transcriptional and phenotypic profiling of brain T cells. **A**, enumeration of extravascular CD8^+^ T cells in 4 month-,10 month-, and 15 month-old WT and APP/PS1dE9 mice by flow cytometry. **B**, frequencies and numbers of TCF-1^+^ CD69^-^, TCF-1^+^ CD69^+^, and TCF-1^-^ CD69^+^ CD8^+^ T cells in the brain of 10 month-old sex-matched WT and APP/PS1dE9 mice; n= 3-6 mice per group. **C**, frequencies and numbers of TCF-1^+^ CD69^-^, TCF-1^+^ CD69^+^, and TCF-1^-^ CD69^+^ CD8^+^ T cells in the brain of 15 month-old sex-matched WT and APP/PS1dE9 mice; n= 3-4 mice per group. **D**, Uniform Manifold Approximation and Projection (UMAP) of scRNA-seq of extravascular brain CD3^+^ T cells from 10 month-old WT and APP/PS1dE9 mice (left), split by genotype (right); n = 3-4 mice per genotype. Cells were processed using the Smart-seq2 protocol (n = 1326 cells). **E,** heatmap of top differentially expressed (DE) genes defining each cluster. **F**, barplot showing the number of cells per T cell cluster in 10 month-old WT vs. APP/PS1dE9 mice. **G**, barplot showing the number of DE genes in a pairwise comparison between WT and APP/PS1dE9 mice per T cell cluster. **H**, frequencies of TNF^+^ IFNγ^+^ and total IFNγ^+^ CD8^+^ T cells in the brain of 15 month-old sex-matched WT and APP/PS1dE9 mice following *ex vivo* stimulation using PMA/ionomycin (left), and MFI of TNF and IFNγ in TNF^+^ IFNγ^+^ CD8^+^ T cells (right). **I,** flow cytometric analysis of PD-1 and TCF-1 co-expression in CD69^+^ CD8^+^ T cells in human brain; n = 5 frontotemproal dementia (FTD) patients. **J**, UMAP plot of snRNA-seq analysis of human brain T cells (n = 1391 cells) derived from the ROSMAP study published by Mathys et al^54^. **K**, frequencies of the T cell clusters in healthy subjects/early AD, intermediate AD, and late AD. Data are representative (B,C,H) or pooled (A) from two independent experiments. Statistical analyses were performed using unpaired two-tailed Student’s t test (A) or two-way ANOVA with Šídák’s multiple comparisons test (B,C,H). MFI; median fluorescence intensity; WT, wild type.

Finally, we asked whether the tissue-imprinted transcriptional makeup of brain CD8^+^ Trm cells also applies to human pathology. Using post-mortem brain samples of patients with frontotemporal dementia, we first confirmed PD-1 and TCF-1 expression by brain CD69^+^ CD8^+^ T cells by flow cytometry (**Figure 3I**). Next, we leveraged a single-nucleus RNA-sequencing (snRNA-seq) dataset of post-mortem human prefrontal cortex from healthy subjects and AD patients with varying extent of neuropathology.^54^ Similar to our findings in the mouse brain, we could identify CD8^+^ T cell subsets differentially expressing *TCF7*, *ITGA1*, and *PDCD1* (**Figure 3J** and **Figures S3A** and **S3B**). In contrast to mouse brain CD8^+^ T cells, we identified a cluster marked by high expression of transcription factors (e.g. *ZEB2*) and effector molecules (e.g. *PRF1*) associated with terminal effector T cells (**Figure S3A**). Comparing healthy subjects/early AD to intermediate- and late-stage AD, we observed no substantial difference in the frequencies of the identified T cell clusters (**Figure 3K** and **Figure S3C**), and pairwise comparison of T cells in healthy/early-AD patients to their counterparts in intermediate- and late-stage AD revealed essentially no DE genes (**Figure S3D**).

Overall, our data indicate that in AD and beta-amyloidosis mouse models, brain tissue residency of CD8^+^ T cells is driven in an age- and disease-dependent manner. Furthermore, our results show that Trm cell phenotypes are largely conserved between mouse and human.

### Acute viral infection generates functionally competent bona-fide brain CD8^+^ Trm cells

Consistent with previous studies,^34,55^ our data demonstrate that a subset of brain CD8^+^ Trm cells express PD-1. Conversely, elevated and sustained PD-1 expression is a classical feature of CD8^+^ T cell exhaustion in chronic viral infection and cancer.^56^ Interestingly, *ex vivo* stimulation revealed that steady-state brain PD-1^+^ Trm cells possessed a greater capacity to produce IFNγ than PD-1^-^ Trm cells (**Figure S4A**). Furthermore, compared to their counterparts in spleen and kidney, brain PD-1^+^ CD8^+^ Trm cells comprised higher frequencies of TNF^+^ IFNγ^+^ and total IFNγ^+^ cells (**Figure S4B**), suggesting that PD-1 expression is a feature of brain Trm cell differentiation but not a state of exhaustion in these cells.

To further test whether PD-1 expression by CD8^+^ Trm cells is linked to exhaustion, we compared extravascular brain CD8^+^ Trm cells in naive mice to those in mice with chronic LCMV (clone 13) infection, a well-established model for CD8^+^ T cell exhaustion.^57,58^ In addition, we investigated the brain CD8^+^ T cell compartment after clearance of systemic infection with acute LCMV Armstrong, a robust model for generation of memory CD8^+^ T cells.^20,21^

LCMV clone 13 infection resulted in a substantially larger number of CD8^+^ T cells in the brain in comparison to naive mice or mice after LCMV Armstrong infection (**Figures 4A** and **B**). This increase in numbers applied to both, total CD8^+^ T cells and those specific for the LCMV immunodominant epitope glycoprotein (gp)33-41 (hereafter referred to as gp33) (**Figure 4B**). Exhausted CD8^+^ T cells upregulate additional inhibitory receptors including TIGIT and TIM-3 (ref. 59). Brain CD8^+^ Trm cells in naive and Armstrong mice showed negligible PD-1 and TIM-3 co-expression, while clone 13-infected mice exhibited a substantial PD-1^+^ TIM-3^+^ CD8^+^ T cell population in the brain (**Figure 4C**). Further, the amount of PD-1 expressed per CD8^+^ T cell was significantly higher in the brain of clone 13-infected mice (**Figure 4D**). Similar to splenic CD8^+^ T cells, brain CD8^+^ Trm cells exhibited a superior capacity to co-produce TNF and IFNγ in Armstrong-immune compared to clone 13-infected mice (**Figure 4E**), which was not attributed to a difference in the frequencies of gp33-specific CD8^+^ T cells (**Figure 4B**). Moreover, the amount of TNF or IFNγ made by brain CD8^+^ T cells in Armstrong-immune mice was larger than their counterparts in chronically infected mice (**Figure 4E**).

**Figure 4.**
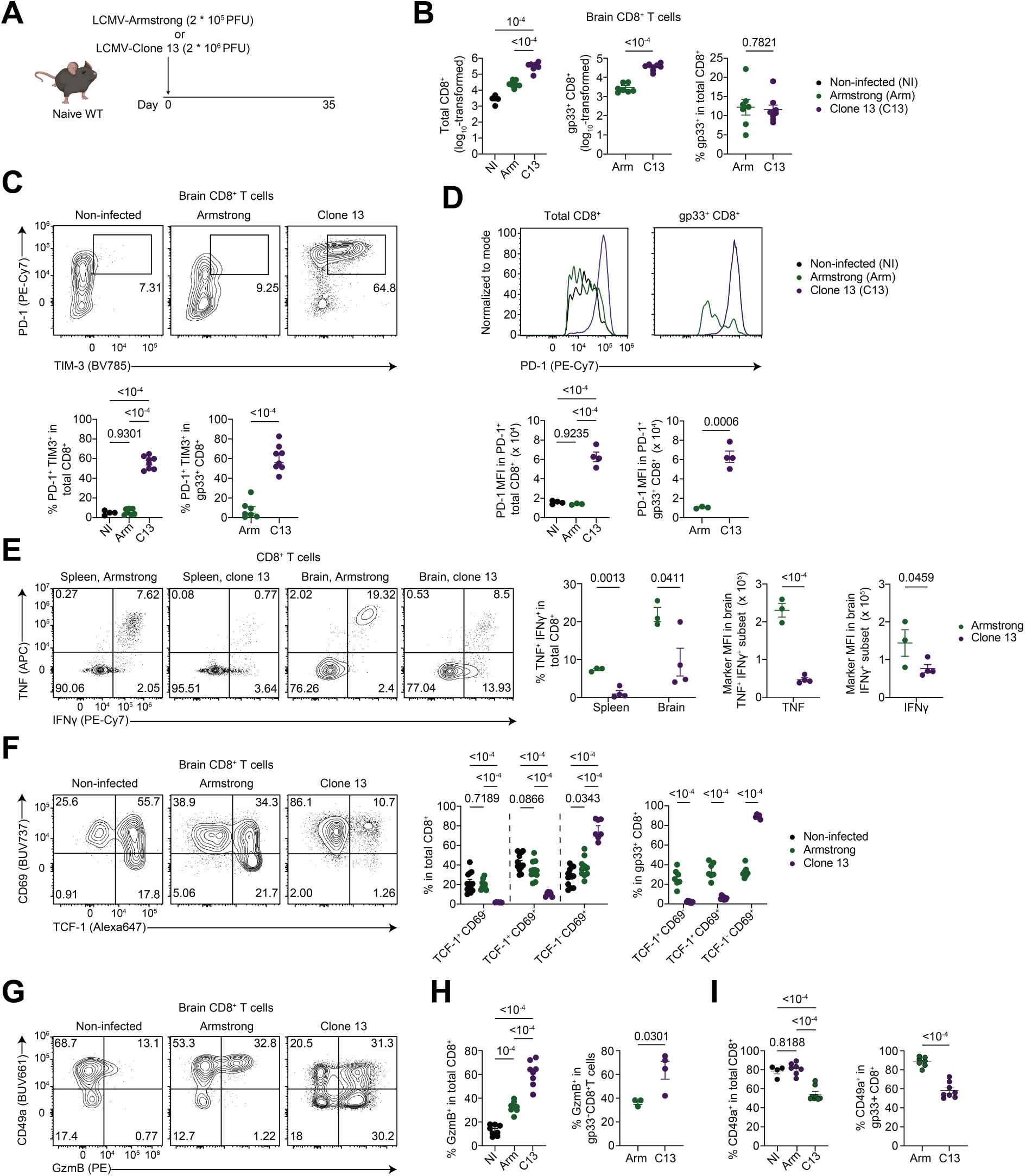
Acute, but not chronic, viral infection generates bona-fide brain CD8^+^ Trm cells. **A**, Nineteen-week-old female mice were either infected with 2 x 10^5^ PFU of LCMV-Armstrong (Arm) or 2 x 10^6^ PFU of LCMV-Clone13 (C13) as indicated, and naive mice were used as controls. **B**, quantification of total CD8^+^ T cells in the brains of naive, Arm, or C13-infected mice, and quantification of numbers and frequencies of glycoprotein 33-41 (gp33)-specific CD8^+^ T cells in Arm and C13-infected mice; n = 4-8 mice per group. **C**, co-expression of PD-1 and TIM3 in brain CD8^+^ T cells in naive, Arm- or C13-infected mice, shown for total and gp33-specific CD8^+^ T cells; n = 4-8 mice per group. **D**, PD-1 MFI in PD-1^+^ brain CD8^+^ T cells, shown for total and gp33-specific CD8^+^ T cells; n = 4 mice per group. **E**, frequency of TNF^+^ IFNγ^+^ splenic and brain CD8^+^ T cells upon ex vivo stimulation with gp33 peptide for 4 hr in the presence of brefeldin A; TNF MFI in TNF^+^ IFNγ^+^ brain CD8^+^ T cells; and IFNγ MFI in total IFNγ^+^ brain CD8^+^ T cells; n = 3-4 mice per group. **F**, frequencies and numbers of TCF-1^+^ CD69^-^, TCF-1^+^ CD69^+^, and TCF-1^-^ CD69^+^ in total and gp33-specific brain CD8^+^ T cells; n = 7-8 mice per group. **G-I**, proportions of GzmB^+^ (**H**) and CD49a^+^ (**I**) cells in brain CD8^+^ T cells, shown for total and gp33-specific CD8^+^ T cells; n = 7-8 mice per group. Data are representative (D-E) or pooled (B,C,F-I) from two independent experiments. GzmB, granzyme B; LCMV, lymphocytic choriomeningitis virus; MFI, median fluorescence intensity.

We next asked how acute or chronic viral infection affected the TCF-1^+^ CD69^+^ subset of brain CD8^+^ Trm cells. In Armstrong-immune mice, we observed no significant changes in the frequencies of these subsets compared to naive mice (**Figure 4F**). Conversely, chronic viral infection resulted in a sharp loss of the brain TCF-1^+^ CD8^+^ Trm cells (**Figure 4F**). Consistent with this loss of TCF-1, chronically infected mice showed a larger frequency of GzmB-producing CD8^+^ T cells in the brain compared to non-infected or Armstrong mice (**Figures 4G** and **4H**). Moreover, brain CD8^+^ Trm cells in chronically infected mice comprised a smaller proportion of CD49a^+^ cells compared to Armstrong-immune mice (**Figures 4G** and **4I**).

Taken together, acute viral infection generates a large pool of bona-fide brain CD8^+^ Trm cells, suggesting that it can be employed as a robust model to study the determinants of brain CD8^+^ Trm cell development. Conversely, chronic viral infection represents an example of a context-specific alteration of the tissue-specific signature of brain CD8^+^ Trm cells described so far.

### TCF-1 promotes differentiation and maintains homeostasis of brain CD8^+^ Trm cells

We next aimed to understand what molecules shape the differentiation and function of brain CD8^+^ Trm cells. Previous studies have associated TCF-1 downregulation with the formation of CD103^+^ Trm cells,^28,60^ and reported that repression of TCF-1 in brain CD8^+^ Trm cells was necessary for the induction of neuropathology in the context of autoimmunity.^36^ However, whether TCF-1 shapes the formation of brain CD8^+^ Trm cells in a cell-intrinsic manner remains unknown. To address this gap in knowledge, we first crossed *Tcf7^fl/fl^* mice to *Cd8^Cre^* mice to achieve CD8^+^ T cell-specific deletion of TCF-1 (**Figure 5A**). We infected *Tcf7^fl/fl^Cd8^Cre^* and *Tcf7^fl/fl^* control mice with LCMV Armstrong and assessed the splenic and brain CD8^+^ T cell compartments on day 44 post-infection (p.i.) (**Figure 5A**). We first confirmed the efficient deletion of TCF-1 in splenic and brain CD8^+^ T cells (**Figure 5B** and **Figure S5A**). Consistent with previous studies,^42,43^ TCF-1 deficiency impaired the development of CD8^+^ Tcm cells in the spleen (**Figure S5B**). In the brain, while the frequency of CD69-expressing CD8^+^ T cells was not altered (**Figure 5B**), *Tcf7^fl/fl^Cd8^Cre^* mice exhibited fewer antigen-specific and total CD8^+^ Trm cells (**Figure 5C**), a reduction that was not observed in the spleen (**Figure S5C**). Furthermore, there was an increased frequency of CD69^+^ GzmB^+^ but a substantially reduced fraction of CD69^+^ PD-1^+^ brain CD8^+^ Trm cells in *Tcf7^fl/fl^ Cd8^Cre^* mice compared to control mice (**Figure 5D** and **Figure S5D**).

**Figure 5.**
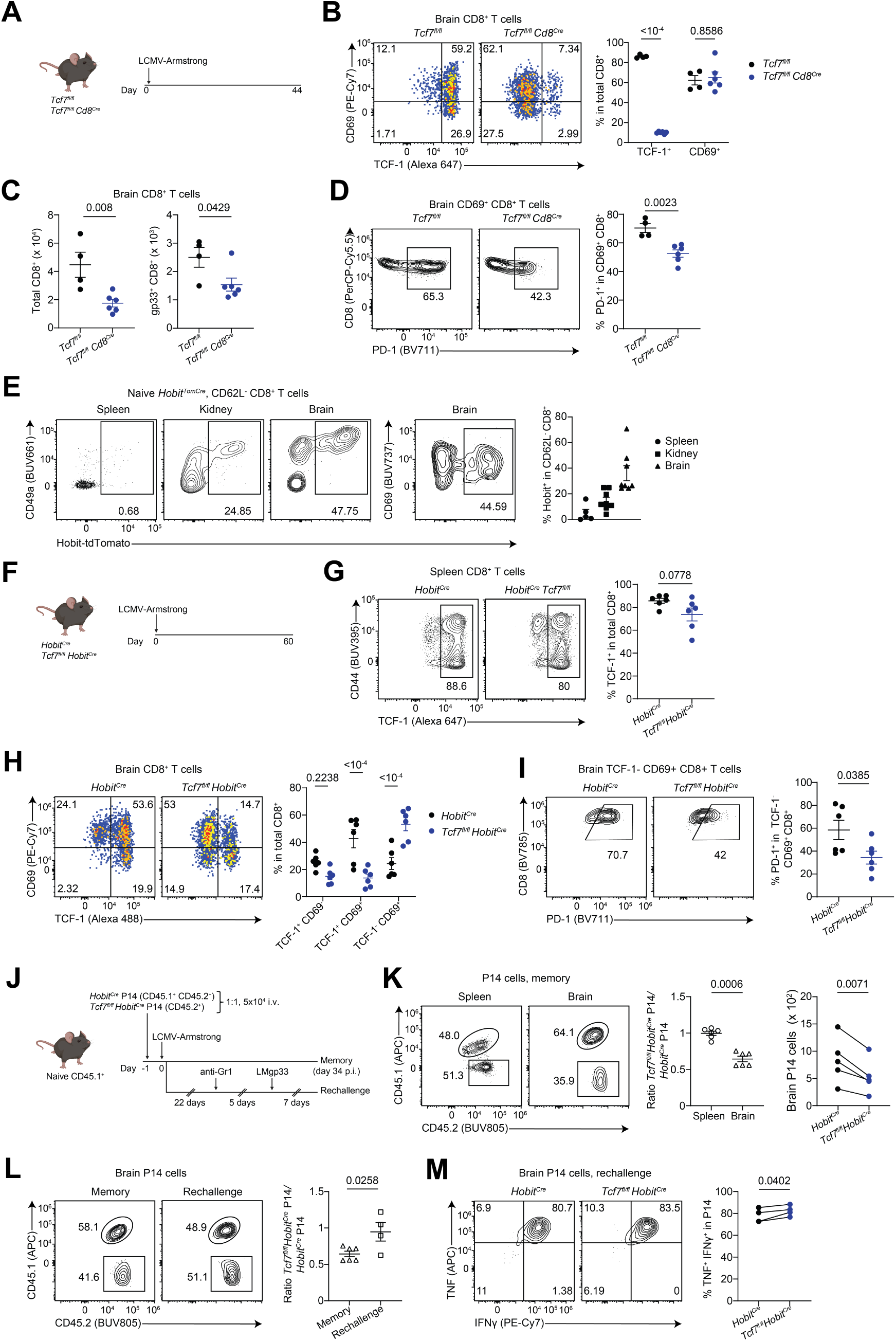
TCF-1 promotes the differentiation and restricts recall of brain CD8^+^ Trm cells. **A**, sex- and age-matched *Tcf7^f/fll^* and *Tcf7^f/fll^ Cd8^Cre^* mice were infected with LCMV-Armstrong (2 x 10^5^ PFU i.p.) and sacrificed on day 44 p.i. **B**, frequencies of TCF-1^+^ and CD69^+^ in total CD8^+^ T cells in the brain; n = 4-6 mice per group. **C**, flow cytometric quantification of total and gp33-specific CD8^+^ T cells in the brain; n = 4-6 mice per group. **D**, proportion of PD-1^+^ in CD69^+^ brain CD8^+^ T cells; n = 4-6 mice per group. **E**, frequencies of Hobit^+^ in splenic CD62L^-^ CD8^+^ and extravascular kidney and brain CD62L^-^ CD8^+^ T cells in naive *Hobit^Cre^* mice, which carries a Cre transgene and tdTomato reporter downstream of *Hobit* promoter; n = 5-8 mice per organ. **F**, sex- and age-matched *Hobit^Cre^* and *Tcf7^fl/fl^ Hobit^Cre^* mice were infected with LCMV-Armstrong (2 x 10^5^ PFU i.p.) and analyzed 60 days p.i. **G**, frequency of TCF-1^+^ in spleen CD8^+^ T cells; n = 6 mice per group. **H**, frequencies of TCF-1^+^ CD69^-^, TCF-1^+^ CD69^+^, and TCF-1^-^ CD69^+^ brain CD8^+^ T cells; n = 6 mice per group. **I**, Percentage of PD-1^+^ in TCF-1^-^ CD69^+^ brain CD8^+^ T cells; n = 6 mice per group. **J**, naive CD45.1 mice received equal numbers of *Hobit^Cre^* P14 (CD45.2) and *Tcf7^fl/fl^ Hobit^Cre^* P14 (CD45.1 CD45.2) cells, which was followed by LCMV-Armstrong infection (2 x 10^5^ PFU) 24 hr later. Mice either were analyzed on day 34 p.i., or received 200 µg of anti-Gr1 i.p. on day 22 p.i., were rechallenged with 1 x 10^5^ CFU of LM-gp33 i.v. 5 days later, and sacrificed 7 days post-rechallenge. **K**, frequencies of *Hobit^Cre^* P14 and *Tcf7^fl/fl^ Hobit^Cre^* P14 in spleen and brain (left), and their respective numbers in brain (right), on day 34 p.i. Flow plots depict CD45.2^+^ CD8^+^ T cells (i.e. total P14); n = 6 mice per genotype. **L**, frequencies of *Hobit^Cre^* P14 and *Tcf7^fl/fl^ Hobit^Cre^* P14 in the brain on day 34 post-Arm vs. day 7 post-rechallenge frequency; n = 4-6 per group. **M**, frequency of TNF^+^ IFNγ^+^ in brain *Hobit^Cre^* P14 and *Tcf7^fl/fl^ Hobit^Cre^* P14 cells on day 7 post-LMgp33 rechallenge, following a 4 hr *ex vivo* stimulation with gp33 in the presence of brefeldin A; n = 4 mice. Data are representative (A-F and L-N) or pooled (H-J) from two independent experiments. Statistical analyses were performed using unpaired (C,D,G,I,L) or paired (K,M) two-tailed Student’s *t* test, or two-way ANOVA with Šídák’s multiple comparisons test (B,H). IFNγ, interferon gamma; i.p., intraperitoneally; LCMV, lymphocytic choriomeningitis virus, PFU, plaque-forming unit; p.i., post-infection; TNF, tumor necrosis factor.

The transcription factor homolog of Blimp1 in T cells (Hobit) is expressed by Trm cells in various tissues.^19^ To investigate the role of TCF-1 in brain CD8^+^ T cells in a Trm cell-specific manner, we utilized a Hobit reporter mouse in which a tdTomato (Tom) cassette and a Cre transgene were inserted into the *Zfp683* (encoding Hobit) locus. Analysis of *Hobit^Cre^* mice revealed that a substantial fraction of brain CD8^+^ Trm cells co-expressed Hobit and the residency markers CD49a and CD69 (**Figure 5E**). Accordingly, we generated *Tcf7^fl/fl^Hobit^Cre^* mice to assess the Trm cell-specific loss of *Tcf7* (**Figure 5F**). On day 60 post-Armstrong infection, the splenic CD8^+^ T cell compartment in *Tcf7^fl/fl^Hobit^Cre^* mice showed no significant reduction of TCF-1^+^ cells and was largely unaltered compared to *Hobit^Cre^* control mice (**Figure 5G** and **Figure S5E**). In contrast, in the brain, there was a marked decrease in the frequency of TCF-1^+^ CD69^+^ CD8^+^ Trm cells in *Tcf7^fl/fl^Hobit^Cre^* mice (**Figure 5H**). Similar to *Tcf7^fl/fl^Cd8^Cre^* mice, TCF-1^-^ CD69^+^ CD8^+^ T cells in *Tcf7^fl/fl^Hobit^Cre^* mice showed a reduced frequency of PD-1^+^ cells compared to controls (**Figure 5I**). To further examine how TCF-1 regulates the differentiation of brain CD8^+^ Trm cells, we co-transferred equal numbers of congenically-marked *Tcf7^fl/fl^Hobit^Cre^* P14 and control *Hobit^Cre^* P14 CD8^+^ T cells that recognize gp33 into congenically distinct host mice, followed by LCMV-Armstrong infection and analysis 34 days later (**Figure 5J**). As expected, *Tcf7^fl/fl^Hobit^Cre^* P14 cells exhibited a ∼50% reduction in TCF-1 expression in the brain but not in the spleen **(Figure S5F**). Importantly, *Tcf7^fl/fl^Hobit^Cre^* P14 cells showed a ∼2-fold numerical deficit in the brain compared to control P14 cells, a reduction that was not observed in the spleen (**Figure 5K**).

We next asked how TCF-1 impacted secondary expansion of brain CD8^+^ Trm cells upon rechallenge. To this end, we used LCMV-immune *Tcf7^fl/fl^Hobit^Cre^* P14 and *Hobit^Cre^* P14 chimeric recipient mice, depleted circulating memory T cells using anti-Gr1 (ref. 61,62), and rechallenged the mice using gp33-expressing *Listeria monocytogenes* (LM-gp33) one week post anti-Gr1 treatment (**Figure 5J)**. Given that LM readily infects the brain,^63,64^ and coupled with the depletion of circulating memory cells via anti-Gr1 treatment, this experimental setup allows for examining Trm cell response to antigenic rechallenge with minimal contribution from circulating memory T cells.^20,62^ Intriguingly, *Tcf7^fl/fl^Hobit^Cre^* P14 cells exhibited a stronger secondary expansion and higher frequencies of TNF^+^ IFNγ^+^ cells compared to control *Hobit^Cre^* P14 cells (**Figures 5L** and **5M**).

Taken together, these data demonstrate that TCF-1 promotes the differentiation of brain CD8^+^ Trm cells while limiting their secondary expansion and effector function.

### PD-1 safeguards the differentiation and recall capacity of brain CD8^+^ Trm cells

There are conflicting reports whether PD-1 promotes or hinders the development of brain CD8^+^ Trm cells.^55,65–67^ Given the impaired PD-1 expression in brain CD8^+^ Trm cells in the absence of TCF-1 (**Figures 5D** and **5I**), we aimed to disentangle the contribution of PD- 1 signaling to the phenotype observed upon conditional *Tcf7* deletion. To better define the role of PD-1 signaling in the differentiation of brain CD8^+^ Trm cells, we generated mixed bone marrow chimeras of *Pdcd1*^-/-^ (PD-1 KO) and congenically marked WT cells at a ratio of 1:1, followed by LCMV Armstrong infection **(Figure 6A**). On day 45-47 p.i, PD-1 KO CD8^+^ T cells outnumbered PD-1-sufficient cells both in the spleen and brain, albeit to a greater extent in the brain (**Figure 6B**). Despite the overall increase of PD-1 KO CD8^+^ T cells, the frequency of gp33-specific CD8^+^ T cells was decreased among PD-1-deficient CD8^+^ T cells compared to their WT counterparts in the brain but not in the spleen (**Figure 6B**). Splenic PD-1 KO CD8^+^ T cells preferentially adopted an effector memory (Tem) phenotype compared to WT CD8^+^ T cells (**Figure S6A**), whereas the frequency of TCF-1^+^ cells was not altered (**Figure S6B**). Conversely, PD-1-deficient CD8^+^ Trm cells in the brain comprised a smaller proportion of TCF-1^+^ cells compared to WT CD8^+^ T cells. (**Figure S6C**). Furthermore, PD-1 KO brain CD8^+^ Trm cells contained fewer CD49a^+^ cells and GzmB^+^ cells relative to their WT counterparts (**Figure S6D** and **S6E**).

**Figure 6.**
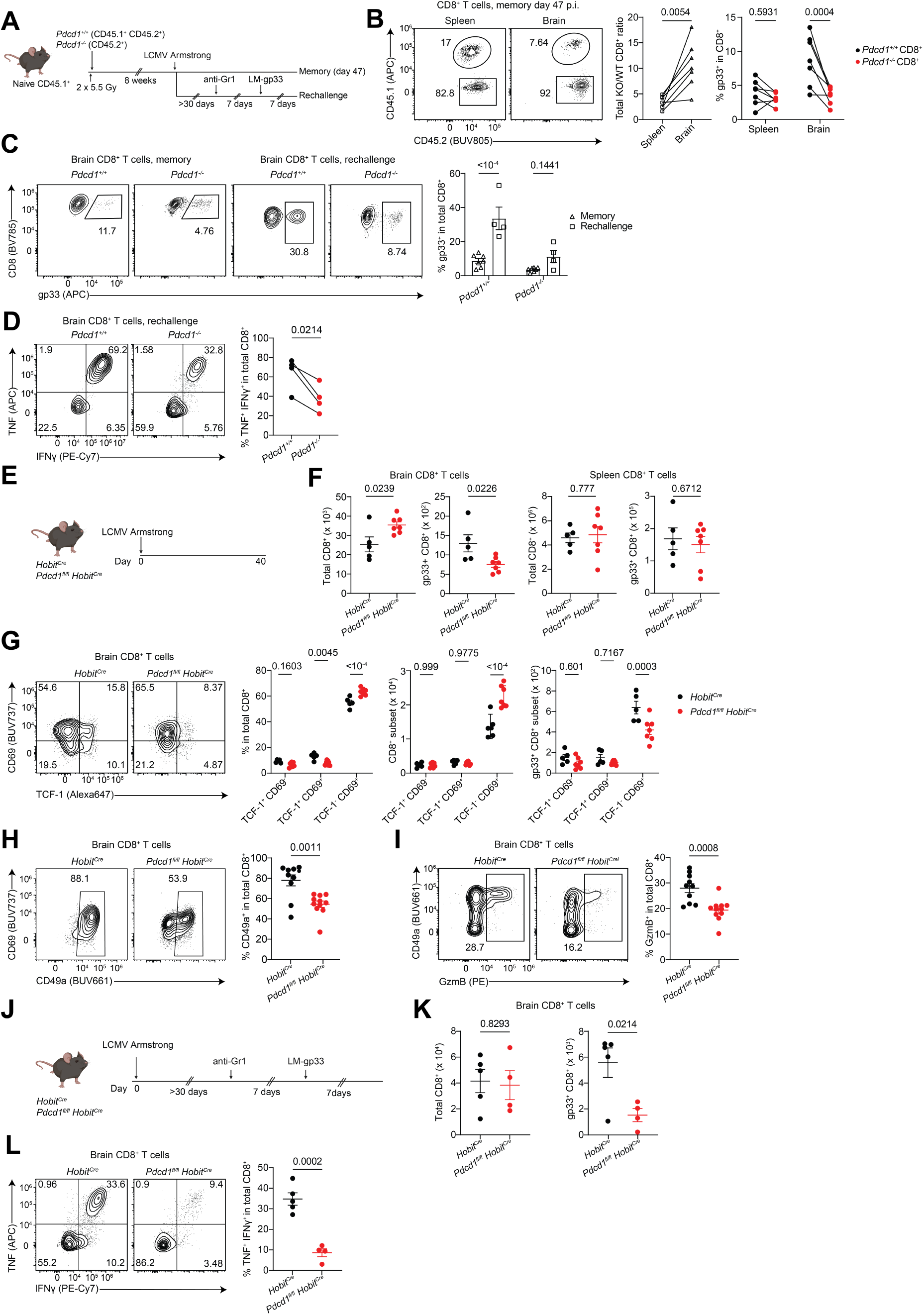
PD-1 maintains brain CD8^+^ T cell phenotype and effector function in a Trm cell-specific manner. **A**, CD45.1 recipient mice were gamma-irradiated twice at 550 Rad followed by bone-marrow reconstitution using hematopoietic stem cells (HSCs) from CD45.1 CD45.2 control and CD45.2 *Pdcd1* KO mice; a total of 4 million HSCs, mixed at a 1:1 ratio from each donor mouse, were administered i.v. to irradiated CD45.1 mice; chimeric mice were infected with LCMV-Armstrong (2 x 10^5^ PFU i.p.), Mice were either analyzed on day 47 p.i., or received > 30 days p.i. 200 µg of anti-Gr1 i.p., and one week later were rechallenged with 1 x 10^5^ CFU of LM-gp33 i.v., and sacrificed 7 days post-rechallenge. **B**, ratio of *Pdcd1* KO to WT CD8^+^ T cells in spleen and brain (left); frequencies of gp33-specific CD8^+^ T cells per genotype in spleen and brain (right); n = 7 mice. **C**, frequency of gp33-specific cells among control and *Pdcd1* KO CD8^+^ T cells during memory and following rechallenge in the brain (c); n = 4-7 mice. **D**, frequency of TNF^+^ IFNγ^+^ in brain CD8^+^ T cells following a 4 hr *ex vivo* stimulation with gp33 in the presence of brefeldin A; n = 4 mice. **E**, sex- and age-matched *Hobit^Cre^* and *Pdcd1^fl/fl^ Hobit^Cre^* mice were infected with LCMV-Armstrong (2 x 10^5^ PFU i.p.) and analyzed 40 days post-infection. **F**, flow cytometric enumeration of total and gp33-specific CD8^+^ T cells in brain and spleen; n = 5-7 mice per group. **G**, frequencies and numbers of TCF-1^+^ CD69^-^, TCF-1^+^ CD69^+^, and TCF-1^-^ CD69^+^ brain CD8^+^ T cells; n = 5-7 mice per group. **H-I**, proportion of CD49a^+^ (**H**) and GzmB^+^ (**I**) among total brain CD8^+^ T cells; n = 10-11 mice per group. **J**, sex- and age-matched *Hobit^Cre^* and *Pdcd1^fl/fl^ Hobit^Cre^* mice were infected with LCMV-Armstrong (2 x 10^5^ PFU i.p.); >30 days later, mice received 200 µg of anti-Gr1 i.p., and one week later mice were rechallenged with 1 x 10^5^ CFU of LM-gp33 i.v., and sacrificed 7 days post-rechallenge. **K**, flow cytometric quantification of total and gp33-specific CD8^+^ T cells in the brain; n = 4-5 mice per genotype. **L**, percentage of TNF^+^ IFNγ^+^ in brain CD8^+^ T cells following 4 hrs *ex vivo* stimulation with gp33 in the presence of brefeldin A; n = 4-5 mice per group. Data are representative (C-G, K-L) or pooled (B, H-I) from 2-3 independent experiments. Statistical analyses were performed using unpaired (F,H,I,K,L) or paired (B left,D) two-tailed Student’s *t* test, or two-way ANOVA with Šídák’s multiple comparisons test (B right,C,G). CFU, colony-forming unit; GzmB, granzyme B; i.p., intraperitoneally; i.v., intravenously; IFNγ, interferon gamma; LCMV, lymphocytic choriomeningitis virus; p.i., post-infection; PFU, plaque-forming unit; TNF, tumor necrosis factor.

Next, we asked how PD-1 modulated the recall response of brain Trm cells upon antigen re-encounter. We treated LCMV-immune PD-1 KO mixed bone marrow chimeras with anti-Gr1 and rechallenged the mice one week later with LM-gp33 (**Figure 6A**). Interestingly, WT gp33-specific CD8^+^ T cells in the brain expanded much better than PD-1 KO cells (**Figure 6C**). In addition, PD-1 KO brain CD8^+^ T cells showed a profound deficit to co-produce TNF and IFNγ relative to their WT counterparts (**Figure 6D**).

To probe whether PD-1 regulated brain CD8^+^ T cells in a Trm cell-specific manner, we generated *Pdcd1^fl/fl^Hobit^Cre^* mice and analyzed LCMV-immune *Pdcd1^fl/fl^Hobit^Cre^* and *Hobit^Cre^* controls on day 40 p.i. (**Figure 6E**). As expected, there was a strong reduction in the fraction of PD-1^+^ brain CD8^+^ T cells in *Pdcd1^fl/fl^Hobit^Cre^* compared to control mice, a reduction that was not observed in the spleen (**Figures S6F** and **S6G**). Consistent with our findings in mixed bone marrow chimeric mice, *Pdcd1^fl/fl^Hobit^Cre^* mice exhibited an increase in total brain CD8^+^ T cell numbers yet a decrease in gp33-specific T cells compared to control mice (**Figure 6F**). Such alterations in brain CD8^+^ Trm cell numbers were exclusively attributed to the TCF-1^-^ subset in *Pdcd1^fl/fl^Hobit^Cre^* mice (**Figure 6G**). Conversely, spleen CD8^+^ T cells showed no difference in total or gp33-specific cell numbers or the representation of naive and memory subpopulations (**Figure 6F** and **Figure S6H**). Furthermore, brain CD8^+^ Trm cells lacking PD-1 contained reduced proportions of CD49a^+^ and GzmB^+^ cells (**Figures 6H** and **6I**). Lastly, we assessed the capacity of CD8^+^ Trm cells lacking PD-1 to mount an effective recall response. We administered anti-Gr1 to LCMV-immune *Pdcd1^fl/fl^Hobit^Cre^* and control mice followed by heterologous rechallenge using LM-gp33 (**Figure 6J)**. Antigen-specific brain CD8^+^ T cells in *Pdcd1^fl/fl^Hobit^Cre^* mice exhibited a marked reduction in their capacity to expand upon antigenic rechallenge compared to *Hobit^Cre^* controls (**Figure 6K**). Further, *Pdcd1^fl/fl^Hobit^Cre^* brain CD8^+^ T cells displayed a dramatic impairment in their ability to produce TNF and IFNγ (**Figure 6L**).

Together, these data indicate that PD-1 signaling is critical for the differentiation and recall capacity of antigen-specific brain CD8^+^ Trm cells.

### TGF-β controls the formation and recall of brain CD8^+^ Trm cells

TGF-β is necessary for the establishment and maintenance of Trm cells in various organs. ^17,21–24^ While previous studies have shown a correlation between the amount of brain Treg cell-derived TGF-β and the formation of brain CD8^+^ Trm cells,^68,69^ it remains unclear whether TGF-β is required for the cell-intrinsic differentiation of brain CD8^+^ Trm cells. We observed that a large proportion of brain CD8^+^ Trm cells expressed CD49a (**Figure S1A**), which is regulated by TGF-β.^26^ Moreover, the majority of brain CD8^+^ Trm cells expressed a TGF-β-induced transcriptional signature^70^ (**Figure S7A**) and *Tgfbr2* (TGF-βRII). To address whether TGF-β controls the differentiation of brain CD8^+^ Trm cells, we crossed *Cd8^Cre^* mice with *Tgfbr2^fl/fl^* mice. We infected *Tgfbr2^fl/fl^Cd8^Cre^* and control (*Tgfbr2^fl/fl^*) mice with LCMV Armstrong and analyzed them on day 60 p.i. (**Figure 7A**). Despite the systemic increase in CD8^+^ Tem cells in *Tgfbr2^fl/fl^Cd8^Cre^* mice (**Figure S7B**), total and gp33-specific brain CD8^+^ Trm cells were significantly decreased compared to controls (**Figure 7B**). In line with the abrogated TGF-β signaling, frequency and numbers of CD49a^+^ CD8^+^ T cells were significantly reduced (**Figure 7C**). In addition, brain CD8^+^ Trm cells in *Tgfbr2^fl/fl^Cd8^Cre^* mice showed modest increases in frequencies of KLRG1 and GzmB expression (**Figures 7D** and **7E**). Taken together, these data demonstrate that cell-intrinsic TGF-β signaling shapes the formation and differentiation of brain CD8^+^ Trm cells.

**Figure 7.**
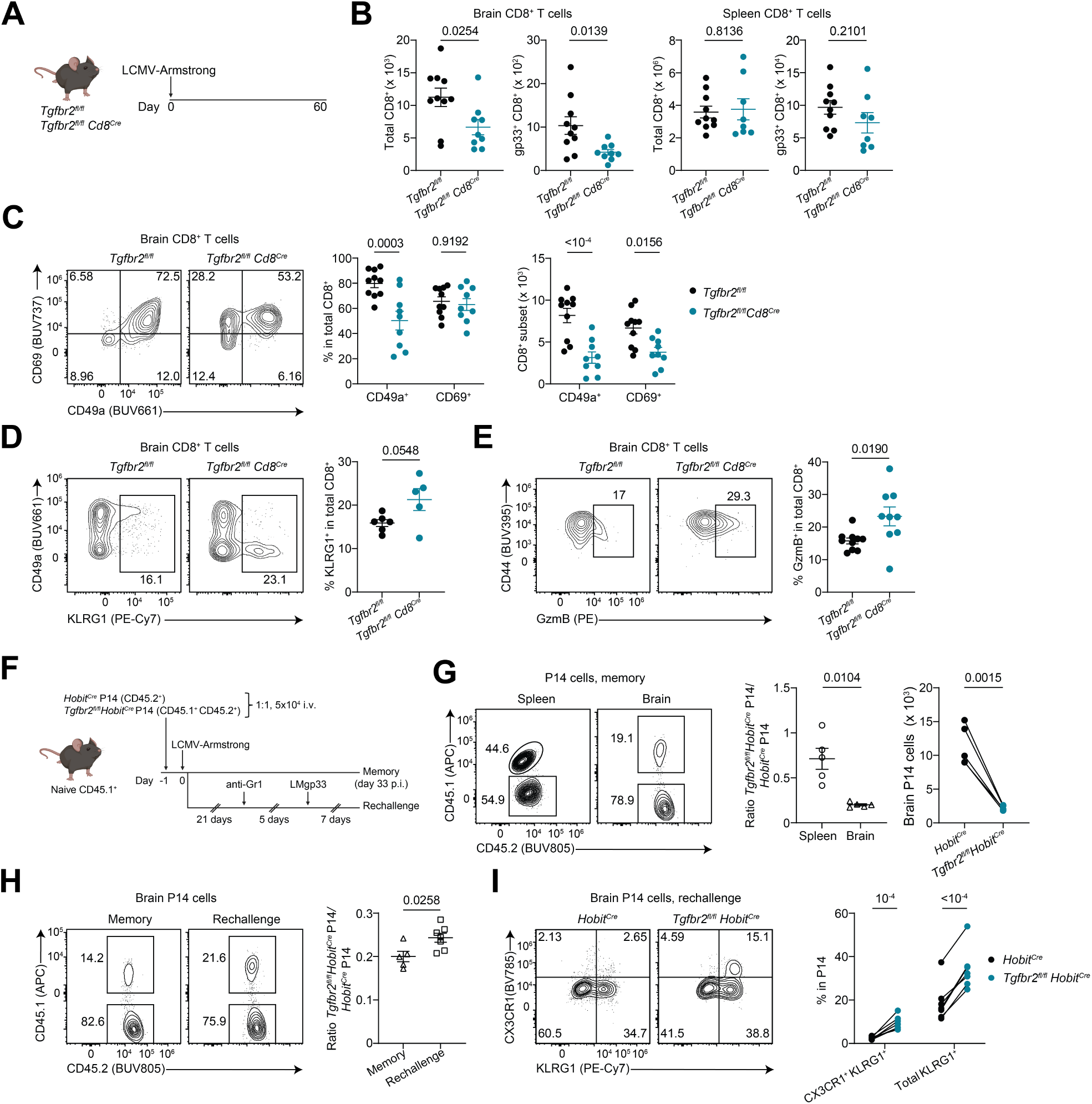
TGF-β signaling is critical for the formation and recall of brain CD8^+^ Trm cells. **A**, sex- and age-matched *Tgfbr2^fl/fl^ Cd8^Cre^* and *Tgfbr2^f/fll^* control mice were infected with LCMV-Armstrong (2 x 10^5^ PFU i.p.) and sacrificed on day 60 post-infection. **B**, flow cytometric quantification of total and gp33-specific CD8^+^ T cells in the brain and spleen; n = 9-10 mice per group. **C**, percentages and numbers of CD49a^+^ and CD69^+^ subsets of brain CD8^+^ T cells; n = 9-10 mice per group. **D**, frequency of KLRG1^+^ in brain CD8^+^ T cells; n = 5-6 mice per group. **E,** proportion of GzmB^+^ in brain CD8^+^ T cells; n = 9-10 mice per group. **F**, naive CD45.1 mice received equal numbers of *Hobit^Cre^* P14 (CD45.2^+^) and *Tgfbr2^fl/fl^ Hobit^Cre^* P14 (CD45.1^+^ CD45.2^+^), which was followed by LCMV-Armstrong infection (2 x 10^5^ PFU) 24hr later. Mice were either analyzed on day 33 p.i., or received 200 µg of anti-Gr1 i.p. on day 21 p.i., were rechallenged with 1 x 10^5^ CFU of LM-gp33 i.v. five days later, and sacrificed 7 days post-rechallenge. **G**, frequencies of *Hobit^Cre^* P14 and *Tgfbr2^fl/fl^ Hobit^Cre^* P14 in spleen and brain (left), and their respective numbers in brain (right), on day 33 p.i. Flow plots depict CD45.2^+^ CD8^+^ T cells (i.e. total P14); n = 5 mice per genotype. **H**, frequencies of *Hobit^Cre^* P14 and *Tgfbr2^fl/fl^ Hobit^Cre^* P14 in the brain on day 33 p.i. vs. day 7 post-rechallenge. Flow plots depict CD45.2^+^ CD8^+^ T cells (i.e. total P14); n = 5-7 mice per condition. **I**, frequencies of CX3CR1^+^ KLRG1^+^ and total KLRG1^+^ in brain *Hobit^Cre^* P14 and *Tgfbr2^fl/fl^ Hobit^Cre^* P14 cells on day 7 post-LM-gp33 rechallenge; n = 7 mice per genotype. Data are representative (D, H-I) or pooled (B,C,E) from 2-3 independent experiments. Statistical analyses were performed using unpaired (B,D,E,H) or paired (G) two-tailed Student’s *t* test, or two-way ANOVA with Šídák’s multiple comparisons test (C,I). CFU, colony-forming unit; GzmB, granzyme B; i.p., intraperitoneally; LCMV, lymphocytic choriomeningitis virus; LM-gp33, gp33-expressing Listeria *monocytogenes*; PFU, plaque-forming unit; p.i., post-infection.

To investigate whether TGF-β modulates brain CD8^+^ T cells in a Trm cell-specific manner, we generated *Tgfbr2^fl/fl^Hobit^Cre^* mice and analyzed LCMV-immune *Tgfbr2^fl/fl^Hobit^Cre^* and control (*Hobit^Cre^*) mice on day 45 p.i. (**Figure S7C**). In the brain, the frequencies of CD49a^+^ CD8^+^ T cells were reduced in *Tgfbr2^fl/fl^Hobit^Cre^* mice compared to controls (**Figure S7D**). In contrast, there were no alterations of splenic CD8^+^ T cells in *Tgfbr2^fl/fl^Hobit^Cre^* mice (**Figure S7E**). In the brain, *Tgfbr2^fl/fl^Hobit^Cre^* mice displayed greater frequencies of GzmB^+^ and KLRG1^+^ brain CD8^+^ Trm cells compared to controls (**Figures S7F** and **S7G**). Moreover, Bcl-2, an anti-apoptotic molecule expressed in CD8^+^ Trm cells,^29^ was downregulated in the absence of TGF-β signaling (**Figure S7H**). To further assess the regulation of brain CD8^+^ Trm cells by TGF-β, we co-transferred *Tgfbr2^fl/fl^Hobit^Cre^* and control P14 cells into congenically marked hosts and analyzed them on day 33 post-Armstrong infection (**Figure 7F**). Intriguingly, *Tgfbr2^fl/fl^Hobit^Cre^* P14 cells in the brain exhibited a ∼4-fold numerical reduction compared controls, a loss that was not observed in the spleen (**Figure 7G**). Moreover, *Tgfbr2^fl/fl^Hobit^Cre^* P14 cells produced less Bcl-2 yet showed greater frequencies of GzmB^+^ cells compared to *Hobit^Cre^* P14 cells (**Figures S7I** and **S7J**).

We next asked how TGF-β impacted the recall response of brain Trm cells. We co-transferred *Tgfbr2^fl/fl^Hobit^Cre^* and control P14 cells into congenically distinct hosts and infected and rechallenged them as described before (**Figure 7F**). Interestingly, brain *Tgfbr2^fl/fl^Hobit^Cre^* P14 cells showed a greater expansion compared to their *Hobit^Cre^* counterparts **(Figure 7H**). In addition, lack of TGF-β signaling facilitated the transition of brain *Tgfbr2^fl/fl^Hobit^Cre^* P14 cells into KLRG1^+^ CX3CR1^+^ (as well as total KLRG1^+^) effector- like cells (**Figure 7I**)

In summary, these data demonstrate that TGF-β plays critical roles in the establishment of brain CD8^+^ Trm cells, the maintenance of their cellular identity, and constraining their transition into effector-like cells upon antigen re-encounter.

## Discussion

Brain Trm cells have been described in physiological and pathophysiological contexts,^37,38,52,71^ yet the molecular landscape of brain CD8^+^ Trm cells at steady state, and to what extent this landscape is altered in disease, remains ill-defined. We report that brain Trm cells adopt a molecular profile primarily instructed by tissue microenvironment rather than ageing or disease context. Indeed, our transcriptional and phenotypic analyses uncovered the same subsets of CD8^+^ T cells in young and aged naive mice, in beta-amyloidosis and AD, and following an acute viral infection. Brain CD8^+^ Trm cells could be classified into distinct subsets based on the combinatorial expression of CD69, TCF-1, PD-1, and Ly6C. Using our novel *Hobit^Cre^* model, we could demonstrate for the first time critical roles for TGFβ signaling and TCF-1 in governing the differentiation of brain CD8^+^ Trm cells. Moreover, we show that PD-1 is necessary for their differentiation and recall in a Trm cell-specific manner.

Tissue-specific imprinting of CD8^+^ Trm cells has been shown for diverse peripheral non-lymphoid tissues. For example, high-dimensional analysis of CD8^+^ Trm cells isolated from various tissues revealed that the tissue microenvironment is a predominant factor in shaping Trm cell transcriptome and phenotype.^20,21^ On the other hand, priming of CD8^+^ T cells with diverse viral and bacterial pathogens resulted in phenotypically similar CD8^+^ Trm cells in the same tissue,^62^ including the brain.^31^ Consistent with our findings, Groh et al reported that CD8^+^ T cells in the aged brain displayed minimal DEGs compared to the same cell types in young mice.^33^ Moreover, bulk transcriptomic profiling of brain CD8^+^ T cells from APP/PS1dE9 mice revealed a substantial transcriptional overlap with their counterparts in WT mice.^72^ Whereas the data presented above support the notion that brain T cell residency instructs a common molecular program, it has long been known that acute intracranial infection generates brain CD8^+^ Trm cells comprising a large fraction of CD103^+^ cells.^27,28^ Conversely, brain CD8^+^ Trm cells generated by a systemic acute infection or through sterile inflammation (e.g. ageing and beta-amyloidosis) consist of a small CD103^+^ population.^31,34,39,72,73^ Notably, systemic chronic viral infection induced a state of exhaustion in brain CD8^+^ T cells, similar to what is described in peripheral tissues.^30,74,75^ Importantly, a recent study has shown that the exhausted T cell fate is imprinted by day 5 post T cell-priming,^59^ suggesting that effector T cells seeding the brain are poised to acquire an exhausted phenotype before tissue entry.

The transcription factor Hobit is expressed in the majority of Trm cells and shapes the differentiation of Trm cells in various tissues.^19,76^ We made use of a *Hobit^Cre^* mouse line to demonstrate Hobit expression by brain CD8^+^ Trm cells and manipulate their gene expression in a Trm cell-specific manner. To our knowledge, this is the first study that employs a Trm cell-specific model to investigate brain CD8^+^ Trm cell differentiation. Importantly, *Hobit^Cre^*-mediated gene deletion did not alter the composition of circulating memory CD8^+^ T cells compared to *Hobit^Cre^* controls, underscoring the utility of the *Hobit^Cre^* model for selective perturbation of Trm cells.

TCF-1 is a transcription factor that plays critical roles in the differentiation and function of central memory CD8^+^ T cells^77^ and marks precursors of memory and exhausted T cells in acute and chronic viral infection.^44,78^ However, few studies have addressed its expression pattern and function in Trm cells.^60,79,80^ We observed a robust fraction of TCF-1^+^ cells among brain CD8^+^ Trm cells in health and disease. Pseudotime analysis of brain CD8^+^ Trm cells inferred a developmental trajectory that progresses along a gradient from a TCF-1^+^ to a TCF-1^-^ cell state, and conditional deletion of *Tcf7* impaired the differentiation of brain CD8^+^ Trm cells. Notably, previous studies have characterized a TCF-1^+^ subset among brain CD8^+^ Trm cells in a model of autoimmune neuroinflammation.^36,81,82^ The authors found that such TCF-1^+^ cells expressed fewer transcripts encoding effector molecules (granzymes and perforin), and that the differentiation into TCF-1^-^ CD8^+^ T cells was required for neuropathology to manifest in mice. The enhanced expression of GzmB by brain TCF-1^-^ CD8^+^ T cells is in agreement with our data. Further, the notion of TCF-1^+^ cells acting as precursors for terminally differentiated brain CD8^+^ T cells^36^ is consistent with our findings. Importantly, we found that brain CD8^+^ Trm cells mounted a stronger recall response in the absence of TCF-1. Accordingly, TCF-1 promotes brain CD8^+^ Trm cell differentiation yet curbs their effector capacity, likely to mitigate collateral tissue damage.

The inhibitory receptor PD-1 is part of a core transcriptional module shared by CD8^+^ Trm cells in diverse tissues in mice and humans.^83,84^ Our data indicate that PD-1 signaling safeguards the differentiation and phenotypic maturation of brain CD8^+^ Trm cells while preserving their functionality and recall capacity. Previous studies employing localized intracranial infection models have addressed how PD-1 regulated brain CD8^+^ Trm cells, with contrasting results.^55,65,67^ Such studies mostly made use of constitutive *Pdcd1* KO (or PD-L1 KO) mice. Given that PD-1 is expressed by additional cell types, including Treg cells, it remains unclear whether PD-1 regulates brain CD8^+^ Trm cells in a cell-intrinsic manner. We addressed this gap in knowledge by generating mixed bone marrow chimera of PD-1 KO and WT cells and generating *Pdcd1^f/fll^ Hobit^Cre^* mice to investigate the Trm cell-intrinsic function of PD-1. Using these models, we could demonstrate a critical role for PD-1 in the differentiation and secondary expansion of brain CD8^+^ Trm cells. Notably, recent evidence points to a similar role for PD-1 in regulating the formation and functional capacity of CD8^+^ Trm cells in the lungs and liver.^85–88^ Accordingly, the timing and impact of PD-1 blockade in humans on brain CD8^+^ Trm cell development will need to be critically evaluated to mitigate the risk of opportunistic infections in the brain.

The cytokine TGF-β is known to be an active player in brain development and homeostasis. Specifically, TGF-β is locally produced in the brain by various cell types, including astrocytes, and regulates neuronal survival and synapse formation.^86^ In the absence of TGF-β signaling, microglia exhibit impaired maturation and compromised long-term survival.^89^ Given the important role exerted by TGF-β in regulating peripheral Trm cells,^23^ coupled with its activity in the brain, we hypothesized that TGF-β signaling may also regulate the formation of CD8^+^ Trm cells. Our data demonstrate a critical role for TGF-β in controlling the establishment and differentiation of CD8^+^ Trm cells in the brain. Furthermore, TGF-β appeared to curb the acquisition of an effector state by brain CD8^+^ Trm cells at the steady state, as well as restraining their capacity to differentiate into effector-like cells upon antigenic rechallenge. Importantly, as small-molecule inhibitors of TGFβ signaling are investigated as therapeutics of lung fibrosis,^90^ the rational design of such molecules is crucial to prevent interference with brain CD8^+^ Trm cell differentiation.

In summary, we found that brain CD8^+^ Trm cells are heterogeneous and adopt a molecular landscape that is strongly driven by the tissue microenvironment but also shaped by the disease in some contexts. We provide evidence on the critical contributions of TCF-1, PD-1, and TGF-β to the formation and function of brain CD8^+^ Trm cells. Accordingly, our study provides a detailed molecular map of brain CD8^+^ Trm cells as well as tools that allow for selectively manipulating them. This lays the foundation for future work to identify and manipulate pathogenic and/or protective subsets of brain CD8^+^ T cells in diverse disease settings.

### Limitations of the study

While we show that TCF-1, PD-1, and TGF-β control the differentiation of brain CD8^+^ Trm cells, our experiments did not disentangle the contributions of such molecules to the establishment and maintenance phases of brain Trm cells. In addition, as it has been reported that glial cells are the main responders to brain CD8^+^ T cells,^91^ it will be interesting to study whether the augmented effector response of brain CD8^+^ Trm cells in the absence of TCF-1 impacts on glial responses. It will also be important to further decipher the differentiation trajectory that CD8^+^ Trm precursors undertake to eventually adopt a mature brain Trm phenotype. Is there an association between a progressive engagement of TCR signaling and brain CD8^+^ T cells differentiation into TCF-1^-^ PD-1^+^ cells? Does TCR signaling contribute to this maturation? Conditional deletion of the TCR on mature or precursors of Trm cells would address this notion. Further, Ly6C is expressed by a large fraction of circulating memory T cells,^17,23^ and its expression was found to be gradually lost in our trajectory inference data. Further investigations could assess if Ly6C^+^ CD69^+^ CD8^+^ T cells in the brain represent early emigrants, and whether Ly6C *per se* is important for the extravasation and infiltration of CD8^+^ T cells into the brain parenchyma.^92^

## RESOURCE AVAILABILITY

### Lead Contact

Further information and requests for resources and reagents should be directed to and will be fulfilled by the Lead Contact, PD Dr. Marc Beyer

### Materials Availability

This study did not generate new unique reagents.

### Data and Code Availability

Raw FASTQ files and processed gene-expression matrices counts for single-cell RNA-sequencing are deposited in the Gene Expression Omnibus under accession number GSE294065. Any additional information required to reanalyze the data reported in this paper is available from the lead contact upon request.

## ACKNOWLEDGEMENTS

We thank Heidi Theis, Andrea Baral, Stephanie Weber, Svenja Bourry, and Dina Hüsson for technical assistance. We would like to thank the Melbourne Cytometry Facility, the Animal Facilities in Bonn and Melbourne for providing support and instrumentation. MDB is supported by the Helmholtz Association and the German Research Foundation (DFG) (SFB1454 project number 432325352, IGK2168/2 project number 272482170). MDB, LB, and ZA are members of the excellence cluster ImmunoSensation2 (EXC2151 project number 390873048). DHDG received funding from the Australian NHMRC (GNT2029937). LB is supported by DFG-funded project ImmuDiet (project number 513977171) and the European Research Council (ERC) under the European Union’s Horizon 2020 research and innovation program (project number 101163024, POLIS). DH was funded by a stipend from the Schlumberger Foundation Faculty of the Future program. DM and LH acknowledge the University of Melbourne for providing a Melbourne Research Scholarship to support the conduct of this study.

The results published here are in part based on data obtained from the AD Knowledge Portal. Raw sequencing data are available via the AD Knowledge Portal (https://adknowledgeportal.org). The AD Knowledge Portal is a platform for accessing data, analyses, and tools generated by the Accelerating Medicines Partnership (AMP-AD) Target Discovery Program and other National Institute on Aging (NIA)-supported programs to enable open-science practices and accelerate translational learning. The data, analyses and tools are shared early in the research cycle without a publication embargo on secondary use. Data is available for general research use according to the following requirements for data access and data attribution (https://adknowledgeportal.synapse.org/Data%20Access). For access to content described in this manuscript see: https://www.synapse.org/Synapse:syn52293417

## AUTHOR CONTRIBUTIONS

Conceptualization: T.E., A.K., and M.D.B; Investigation: T.E., C.S., M.H.S, D.H., D.M., M.K., T.M., Y.L., L.H., C.O.S, and K.M.; Data curation and formal analysis: T.E., A.F., R.S, J.S.S., L.B., K.H., and E.D.D.; Resources: V.B., A.H., D.G., M.F., and Z.A.; Supervision and funding acquisition: A.K., K.M., and M.D.B.; Visualization: T.E., M.D.B.; Writing – original draft: T.E., K.M., A.K., and M.D.B.; writing – editing: all authors.

---

**Figure S1.**
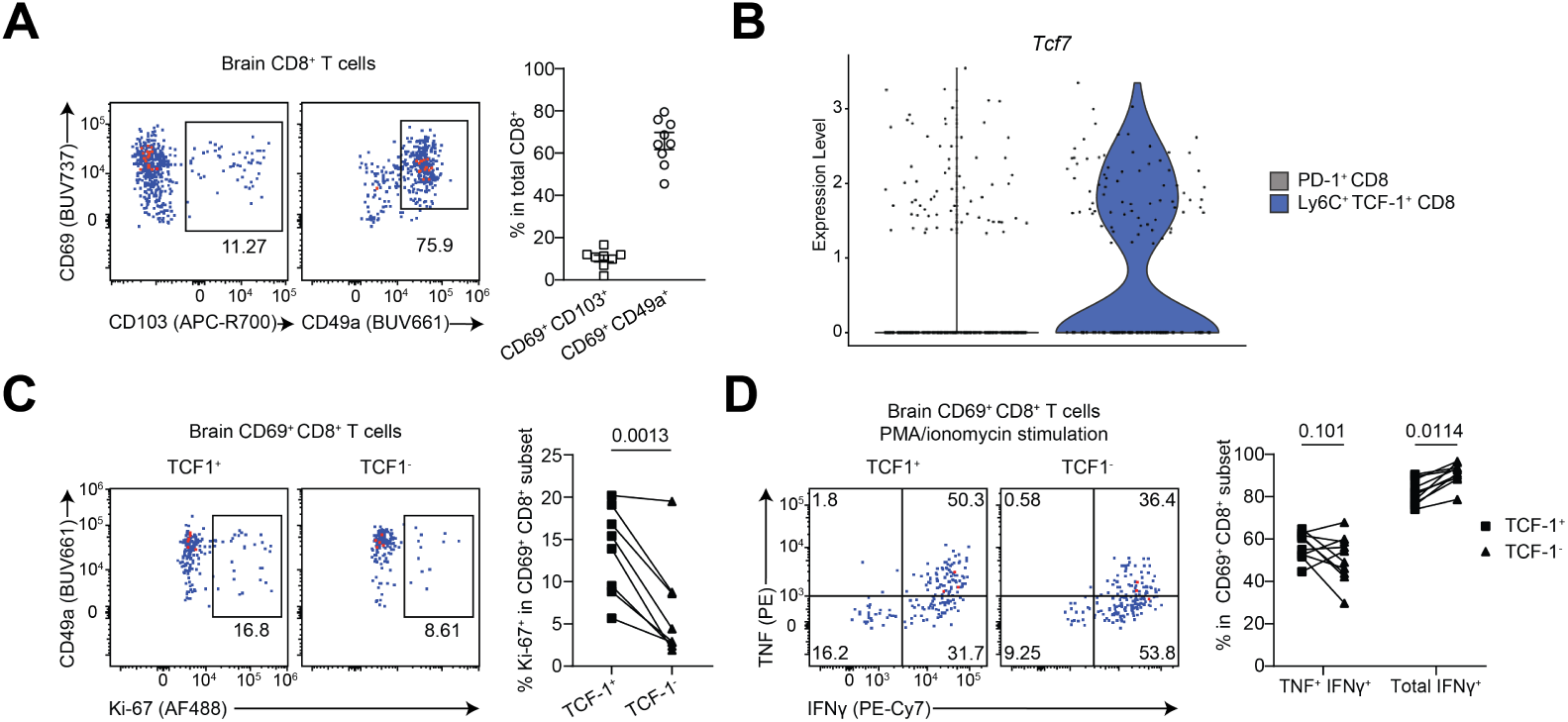
Phenotype and effector function of brain CD8^+^ T cells at steady state (related to Figure 1). **A**, frequencies of CD69^+^ CD103^+^ and CD69^+^ CD49a^+^ subsets of brain CD8^+^ T cells; n = 8 mice. **B**, scRNA-seq violin plot showing *Tcf7* RNA expression in brain CD8^+^ T cells at steady state. **C**, frequency of Ki-67^+^ among TCF-1^+^ and TCF-1^-^ subsets of brain CD69^+^ CD8^+^ T cells; n = 8 mice. **D**, proportions of TNF^+^ IFNγ^+^ and total IFNγ^+^ in TCF-1^+^ and TCF-1^-^ subsets of brain CD69^+^ CD8^+^ T cells following *ex vivo* stimulation with PMA/ionomycin for 4 hr in the presence of brefeldin A; n = 8 mice. Data are pooled from 2-3 independent experiments. Statistical analyses were performed using paired two-tailed Student’s *t* test (C) or two-way ANOVA with Šídák’s multiple comparisons test (D).

**Figure S2.**
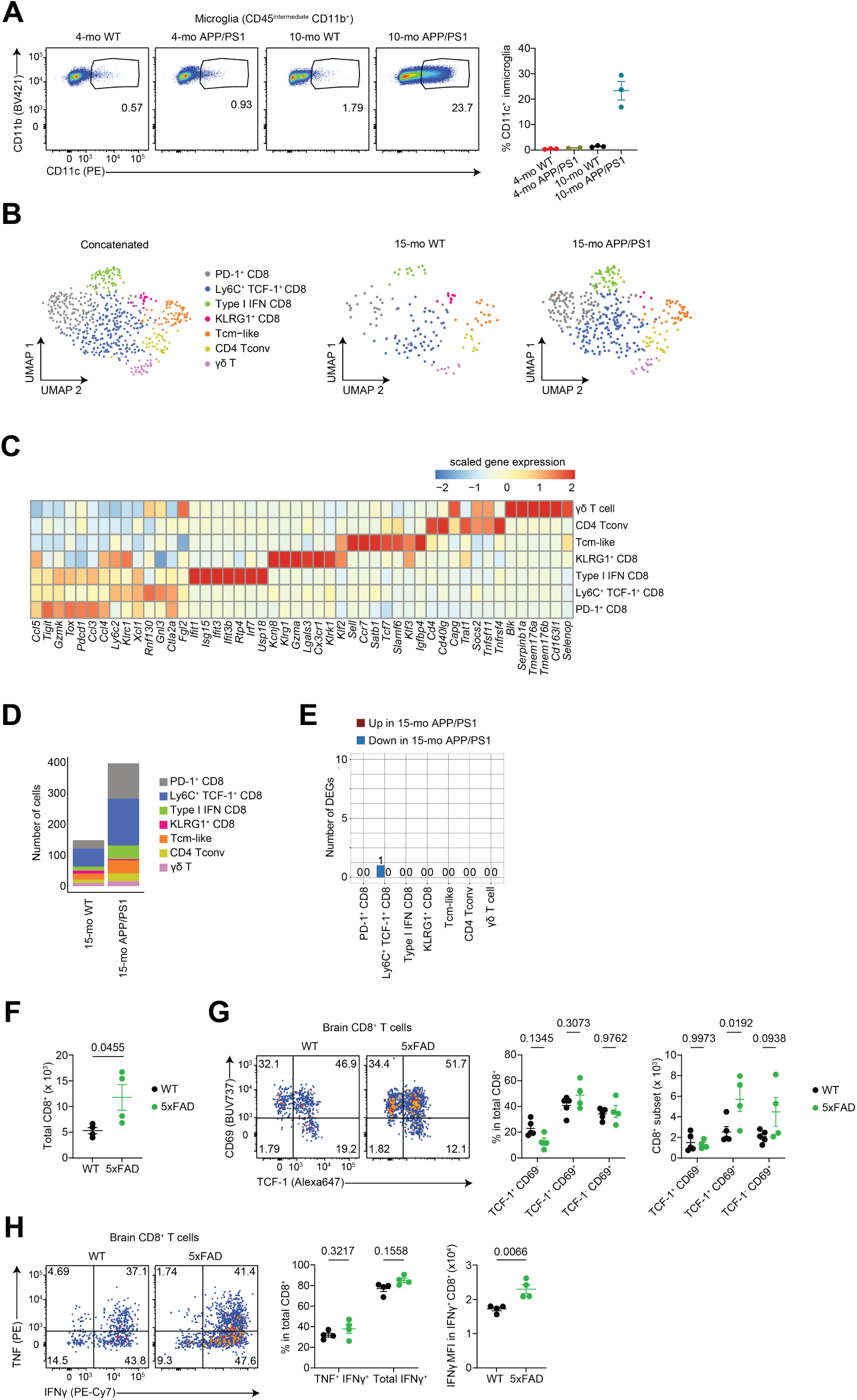
Limited transcriptional and functional alterations of CD8^+^ T cells in the context of beta-amyloidosis (related to Figure 2). **A**, frequency of CD11c^+^ in microglia; 2-3 mice per group. **B**, scRNA-seq of extravascular brain CD3^+^ T cells from 15 month-old WT and APP/PS1dE9 mice; n = 3-4 mice per group. Cells were processed using the FLASH-seq protocol. **C**, heatmap of top differentially expressed (DE) genes defining each cluster. **D**, barplot showing the number of cells per T cell cluster in 15-month-old WT vs. APP/PS1dE9 mice. **E**, barplot depicting DE genes between WT and APP/PS1dE9 for each T cell cluster. **F**, enumeration of extravascular CD8^+^ T cells in 9-month-old 5xFAD mice and sex- and age-matched control mice; n = 4 mice per group. **G**, frequencies and numbers of TCF-1^+^ CD69^-^, TCF-1^+^ CD69^+^, and TCF- 1^-^ CD69^+^ brain CD8^+^ T cells; n = 4 mice per group. **H**, frequencies of TNF^+^ IFNγ^+^ and total IFNγ^+^ CD8^+^ T cells in the brain of 9-month-old sex-matched WT and 5xFAD mice following *ex vivo* stimulation with PMA/ionomycin (left), and MFI of IFNγ in IFNγ^+^ CD8^+^ T cells (right). Data are representative (A, F-H) from 2-3 independent experiments. Statistical analyses were performed using unpaired two-tailed Student’s *t* test (F,H right) or two-way ANOVA with Šídák’s multiple comparisons test (G,H left). DE, differentially expressed; MFI; median fluorescence intensity; WT, wild type.

**Figure S3.**
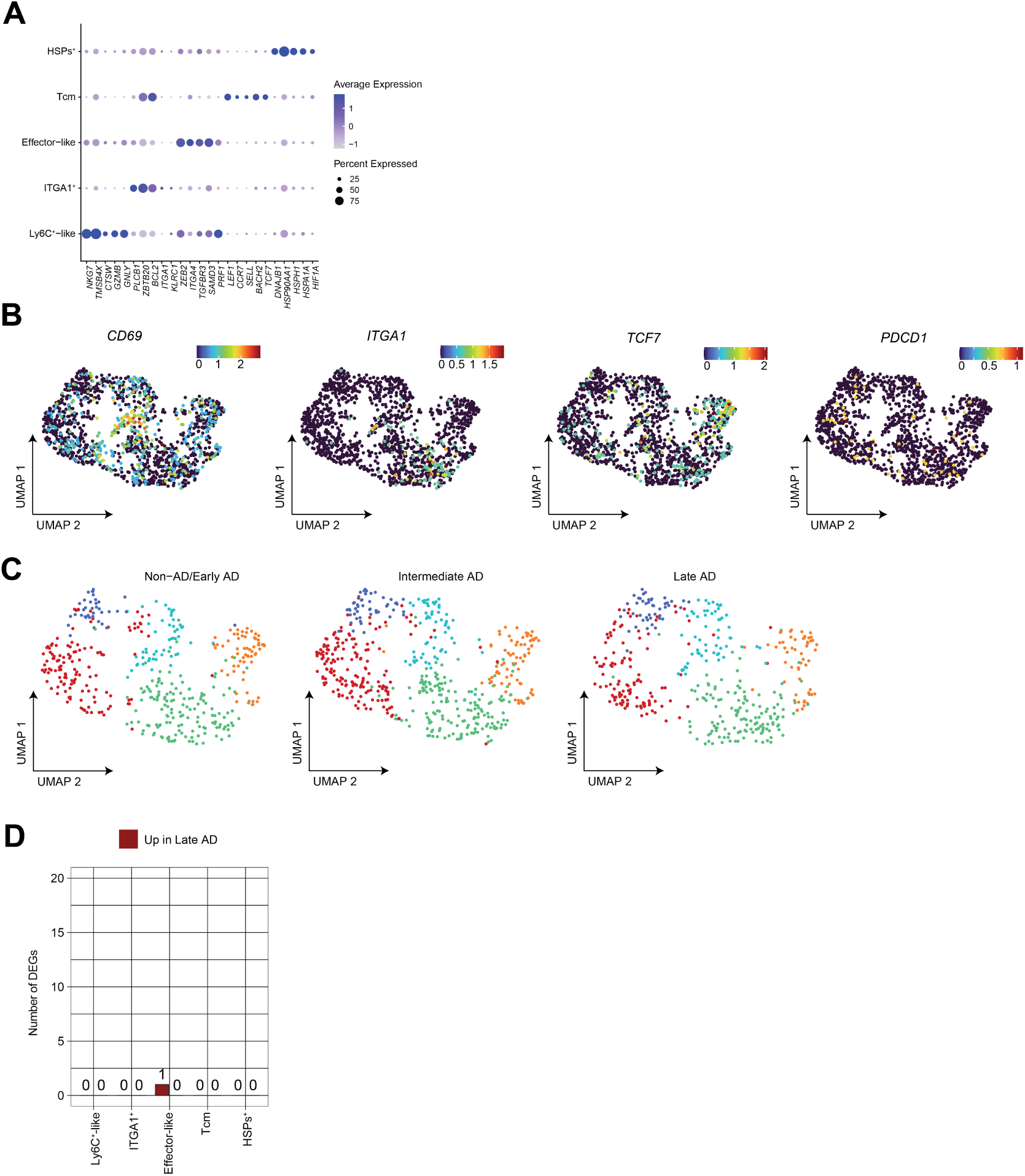
Transcriptional heterogeneity of CD8^+^ T cells in the human brain (related to Figure 3). **A**, dotplot of the top differentially expressed genes defining each T cells cluster. **B**, FeaturePlots of a selected set of genes overlaid on the UMAP described in Figure 3J. **C**, UMAP shown in Figure 3J split by disease stage, namely healthy subjects/early AD, intermediate AD, and late AD. **D**, barplot showing the number of differentially expressed genes between healthy subjects/early AD and late AD per T cell cluster.

**Figure S4.**
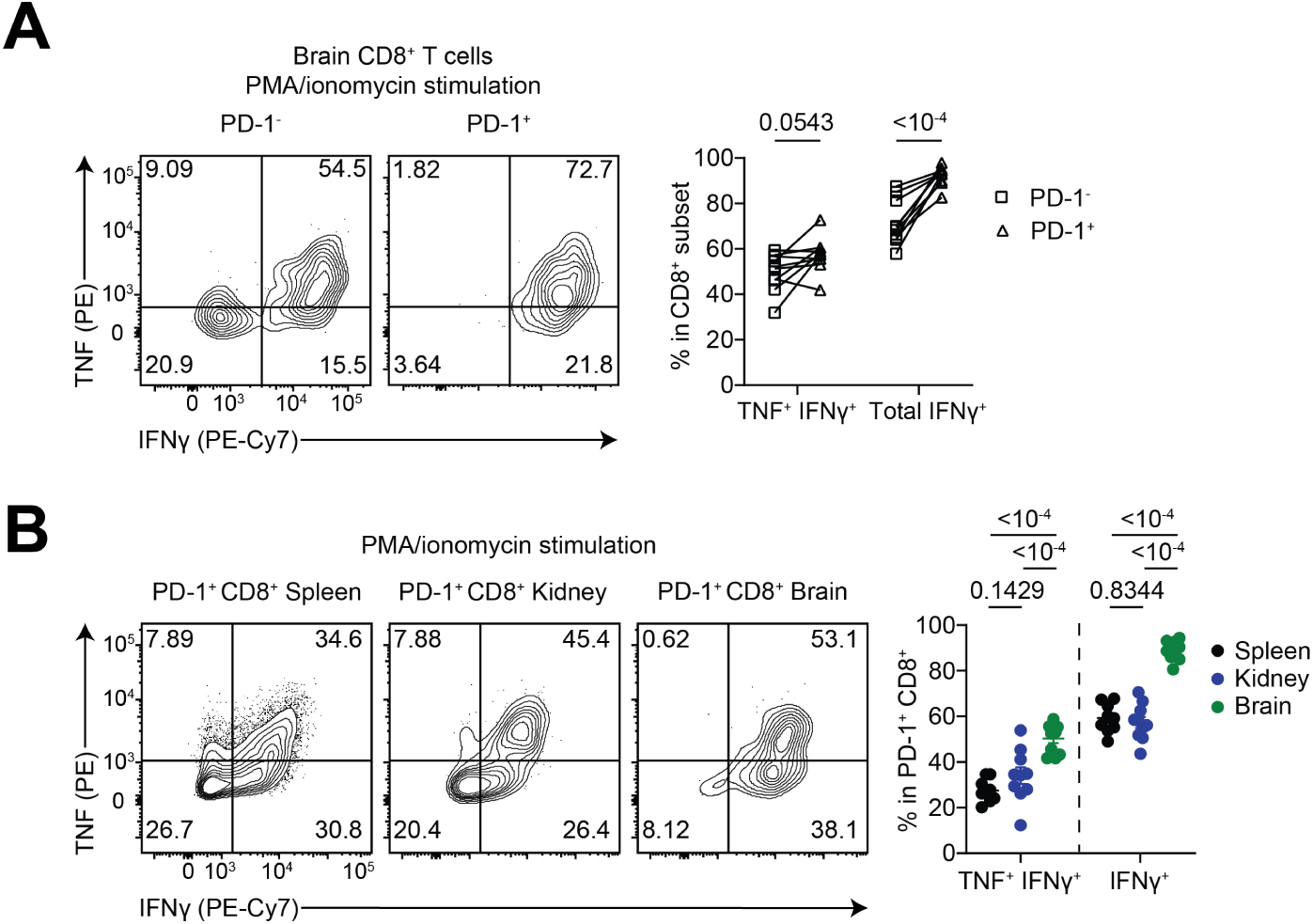
PD-1^+^ CD8^+^ brain Trm cells do not exhibit functional exhaustion. **A**, proportions of TNF^+^ IFNγ^+^ and total IFNγ^+^ in PD-1^-^ and PD-1^+^ brain CD8^+^ T cells following *ex vivo* stimulation with PMA/ionomycin for 4 hr in the presence of brefeldin A; n = 8 mice. **B**, proportions of TNF^+^ IFNγ^+^ and total IFNγ^+^ in PD-1^+^ CD8^+^ T cells in spleen, kidney, and brain following *ex vivo* stimulation with PMA/ionomycin in the presence of brefeldin A; n = 8 mice. Data are pooled from two independent experiments Statistical analyses were performed using two-way ANOVA with Šídák’s multiple comparisons test (A) or Tukey’s multiple comparisons test (B).

**Figure S5.**
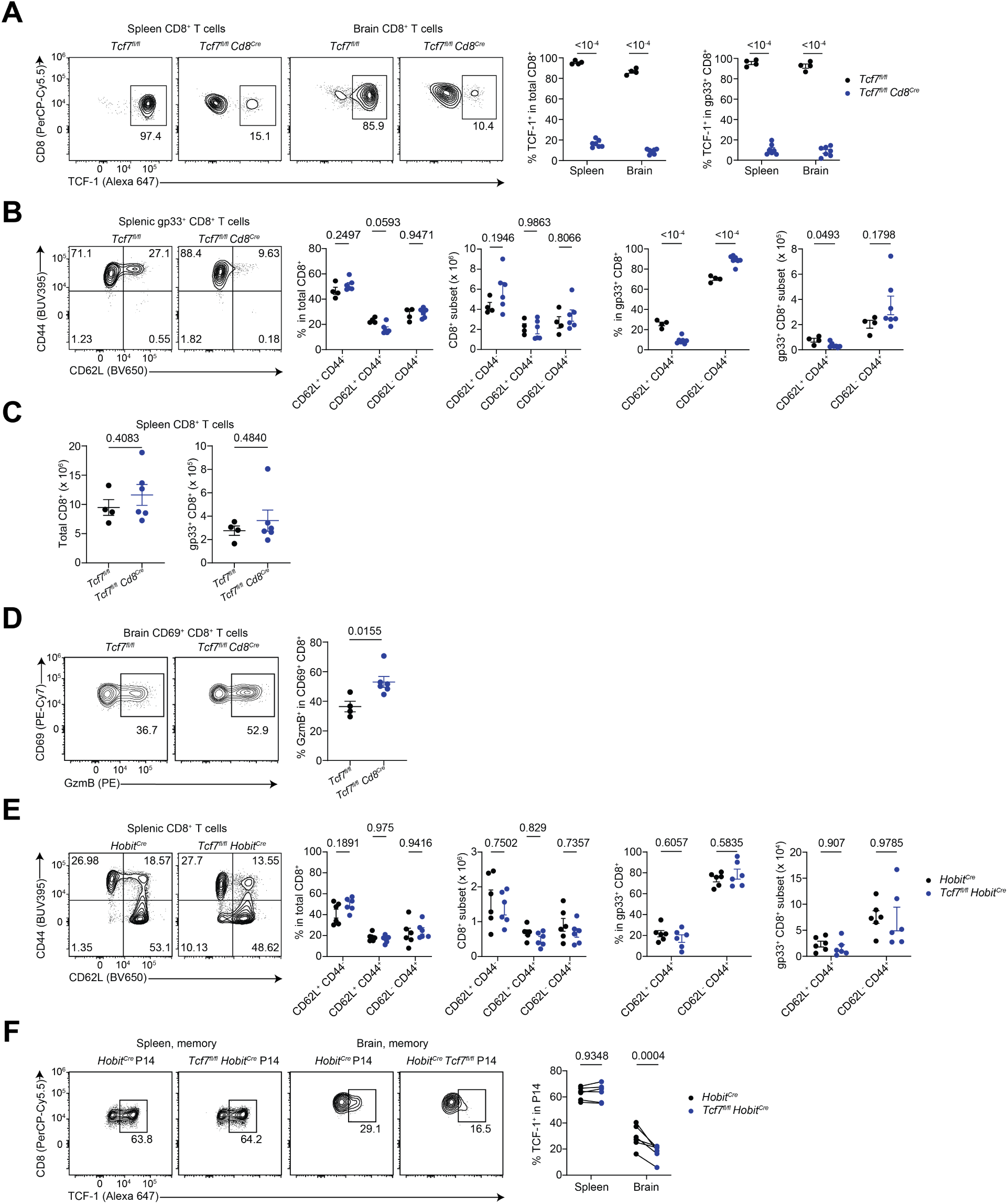
TCF-1 deletion downstream of Hobit does not alter the composition of spleen CD8^+^ T cells (related to Figure 5). **A-B**, sex- and age-matched *Tcf7^f/fll^* and *Tcf7^f/fll^ Cd8^Cre^* mice were infected with LCMV- Armstrong (2 x 10^5^ PFU i.p.) and sacrificed on day 44 p.i. **A**, frequency of TCF-1^+^ in total and gp33-specific CD8^+^ T cells in spleen and brain; n = 4-7 mice per group. **B**, frequencies and numbers of CD62L^+^ CD44^-^, CD62L^+^ CD44^+^, and CD62L^-^ CD44^+^ CD8^+^ T cells in spleen; n = 4-6 mice per group. **C**, flow cytometric quantification of total and gp33-specific CD8^+^ T cells in the brain and spleen; n = 4-6 mice per group. **D**, percentage of GzmB^+^ in brain total CD8^+^ T cells; n = 4-6 mice per group. **E**, sex- and age-matched *Tcf7^fl/fl^ Hobit^Cre^* and *Hobit^Cre^* control mice were infected with LCMV-Armstrong (2 x 10^5^ PFU i.p.) and sacrificed on day 60 p.i. Frequencies and numbers of CD62L^+^ CD44^-^, CD62L^+^ CD44^+^, and CD62L^-^ CD44^+^ CD8^+^ T cells in spleen; n = 6 mice per group. **F**, frequencies of TCF- 1^+^ in *Hobit^Cre^* P14 and *Tcf7^fl/fl^ Hobit^Cre^* P14 in spleen and brain on day 34 post-Armstrong infection; n = 6 mice per genotype. Data are representative (A-D, F) or pooled (E) from two independent experiments. Statistical analyses were performed using unpaired two- tailed Student’s *t* test (C,D) or two-way ANOVA with Šídák’s multiple comparisons test (A,B,E,F). i.p., intraperitoneally; LCMV, lymphocytic choriomeningitis virus, PFU, plaque- forming unit; p.i., post-infection.

**Figure S6.**
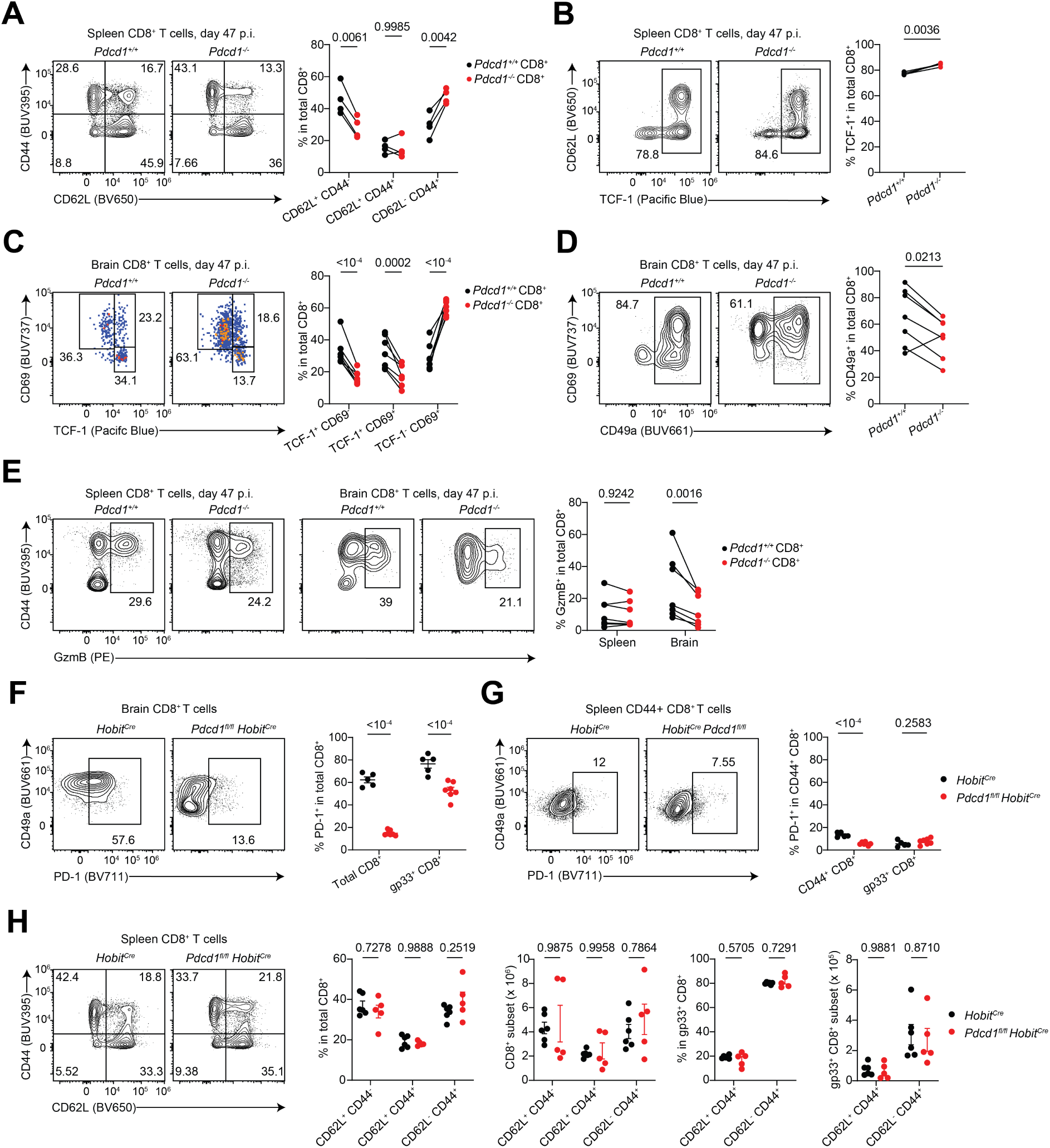
PD-1 promotes the differentiation of brain CD8^+^ Trm cells (related to Figure 6). **A-E**, CD45.1 recipient mice were gamma-irradiated twice at 550 Rad, followed by bone- marrow reconstitution using hematopoietic stem cells (HSCs) from CD45.1 CD45.2 control and CD45.2 *Pdcd1* KO mice; a total of 4 million HSCs, mixed at a 1:1 ratio from each donor mouse, were administered i.v. to irradiated CD45.1 mice; chimeric mice were infected with LCMV-Armstrong (2 x 10^5^ PFU i.p.) and analyzed on day 45-47 p.i. **A**, frequencies of CD62L^+^ CD44^-^, CD62L^+^ CD44^+^, and CD62L^-^ CD44^+^ CD8^+^ T cells in spleen; n = 7 mice. **B**, frequency of TCF-1^+^ in spleen CD8^+^ T cells; n = 3 mice. **C**, proportions of TCF-1^+^ CD69^-^, TCF-1^+^ CD69^+^, and TCF-1^-^ CD69^+^ subsets in brain CD8^+^ T cells; n = 7 mice. **D**, frequency of CD49a^+^ in brain CD8^+^ T cells; n = 7 mice. **E**, proportion of GzmB^+^ in spleen and brain CD8^+^ T cells; n = 7 mice. **F-H**, Sex- and age-matched *Hobit^Cre^* and *Pdcd1^fl/fl^ Hobit^Cre^* mice were infected with LCMV-Armstrong (2 x 10^5^ PFU i.p.) and analyzed 40 days p.i. **F**-**G**, frequencies of PD-1^+^ cells in brain (**F**) and spleen (**G**) CD44^+^ and gp33-specific CD8^+^ T cells; n = 5-7 mice per group. **H**, frequencies and numbers of CD62L^+^ CD44^-^, CD62L^+^ CD44^+^, and CD62L^-^ CD44^+^ CD8^+^ T cells in spleen; n = 5-6 mice per group Data are representative (B, F-G) or pooled (A, C-E, H) from two independent experiments. Statistical analyses were performed using paired two-tailed Student’s *t* test (B,D), or two-way ANOVA with Šídák’s multiple comparisons test (A,C,E-H). GzmB, granzyme B; i.p., intraperitoneally; LCMV, lymphocytic choriomeningitis virus; PFU, plaque-forming unit; p.i., post-infection.

**Figure S7.**
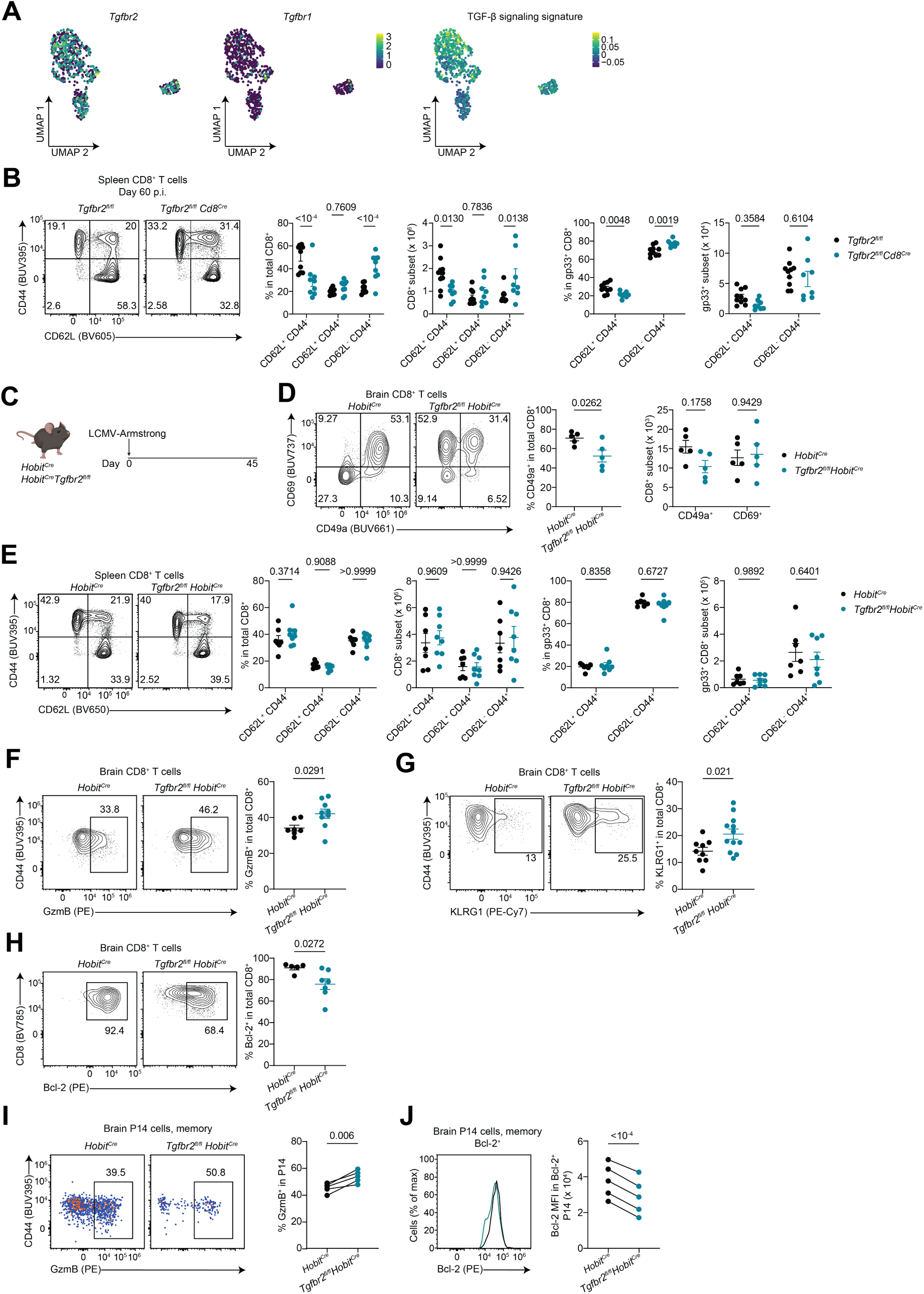
TGF-β signaling promotes the differentiation of brain CD8^+^ T cell residency (related to Figure 7). **A**, scRNA-seq of brain T cells, related to Figure 1. FeaturePlots depicting the expression of *Tgfbr2*, *Tgfbr1*, and gene-expression score of TGF-β signaling, derived from Nath et al^70^. **B**, frequencies and numbers of CD62L^+^ CD44^-^, CD62L^+^ CD44^+^, and CD62L^-^ CD44^+^ among total and gp33-specific CD8^+^ T cells in the spleen on day 60 p.i.; n = 9-10 mice per group. **C**, sex- and age-matched *Hobit^Cre^* and *Tgfbr2^fl/fl^ Hobit^Cre^* mice were infected with LCMV-Armstrong (2 x 10^5^ PFU i.p.) and sacrificed on day 45 p.i. **D**, frequency of CD49a^+^ and absolute numbers of CD49a^+^ and CD69^+^ brain CD8^+^ T cells; n = 5 mice per genotype. **E**, frequencies and numbers of CD62L^+^ CD44^-^, CD62L^+^ CD44^+^, and CD62L^-^ CD44^+^ CD8^+^ T cells in spleen; n = 7-8 mice per genotype. **F**, frequency of GzmB^+^ in brain CD8^+^ T cells; n = 7-9 mice per genotype. **G**, frequency of KLRG1^+^ in brain CD8^+^ T cells; n = 9-12 mice per genotype. **H**, proportion of Bcl-2^+^ in brain CD8^+^ T cells; n = 3-4 mice per group. **I-J**, percentage of GzmB^+^ (**I**) and Bcl-2 MFI in Bcl-2^+^ (**J**) in co-transferred *Hobit^Cre^* P14 and *Tgfbr2^fl/fl^ Hobit^Cre^* P14 cells in the brain of congenically distinct recipients on day 33 p.i.; n = 5 mice. Data are representative (D, I-J) or pooled (B, E-H) from 2-3 independent experiments. Statistical analyses were performed using unpaired (D left, F- H) or paired (I-J) two-tailed Student’s *t* test, or two-way ANOVA with Šídák’s multiple comparisons test (B,D left, E). i.p., intraperitoneally; LCMV, lymphocytic choriomeningitis virus, PFU, plaque-forming unit; p.i., post-infection.

## STAR Methods

### Key Resources Table

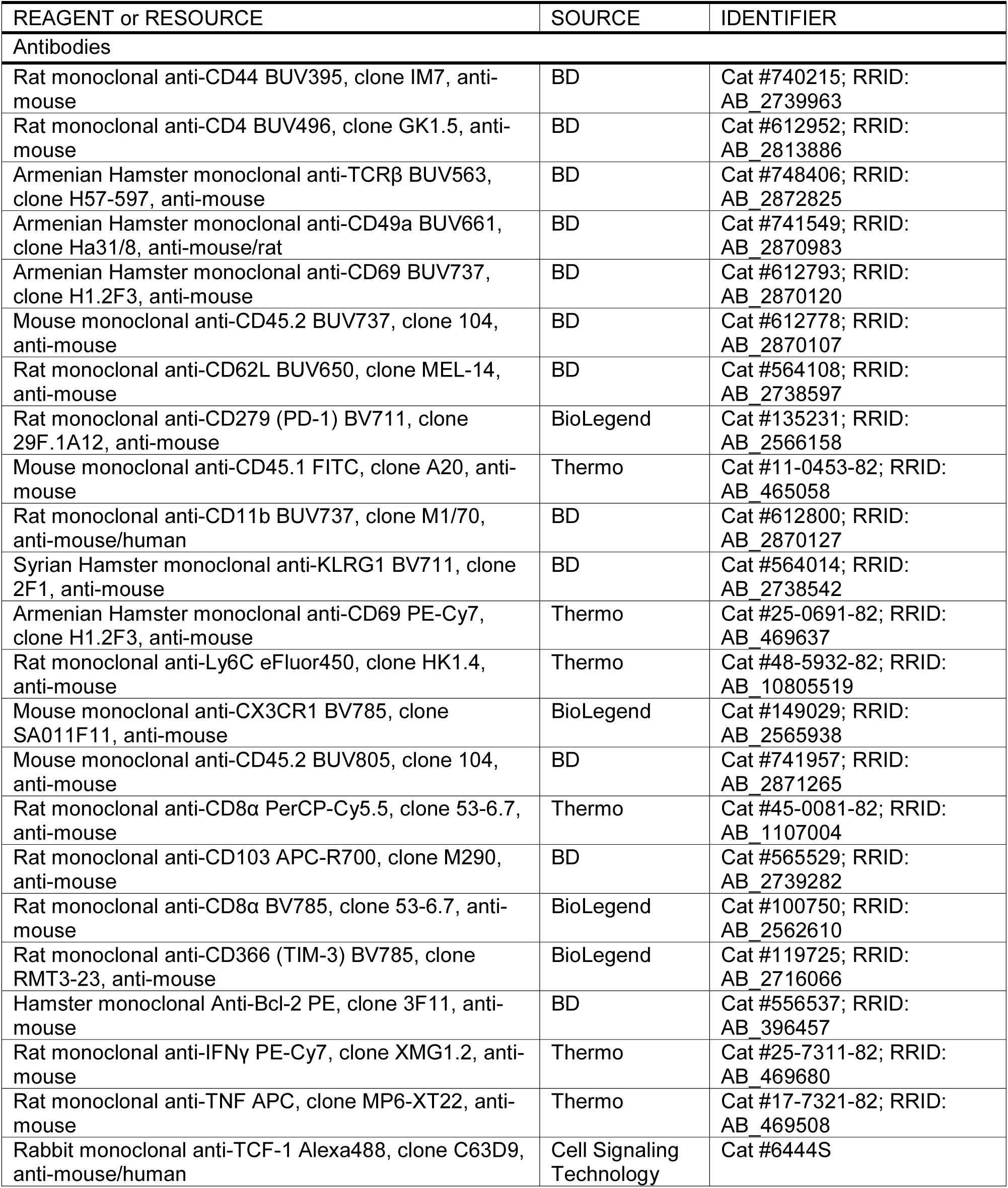
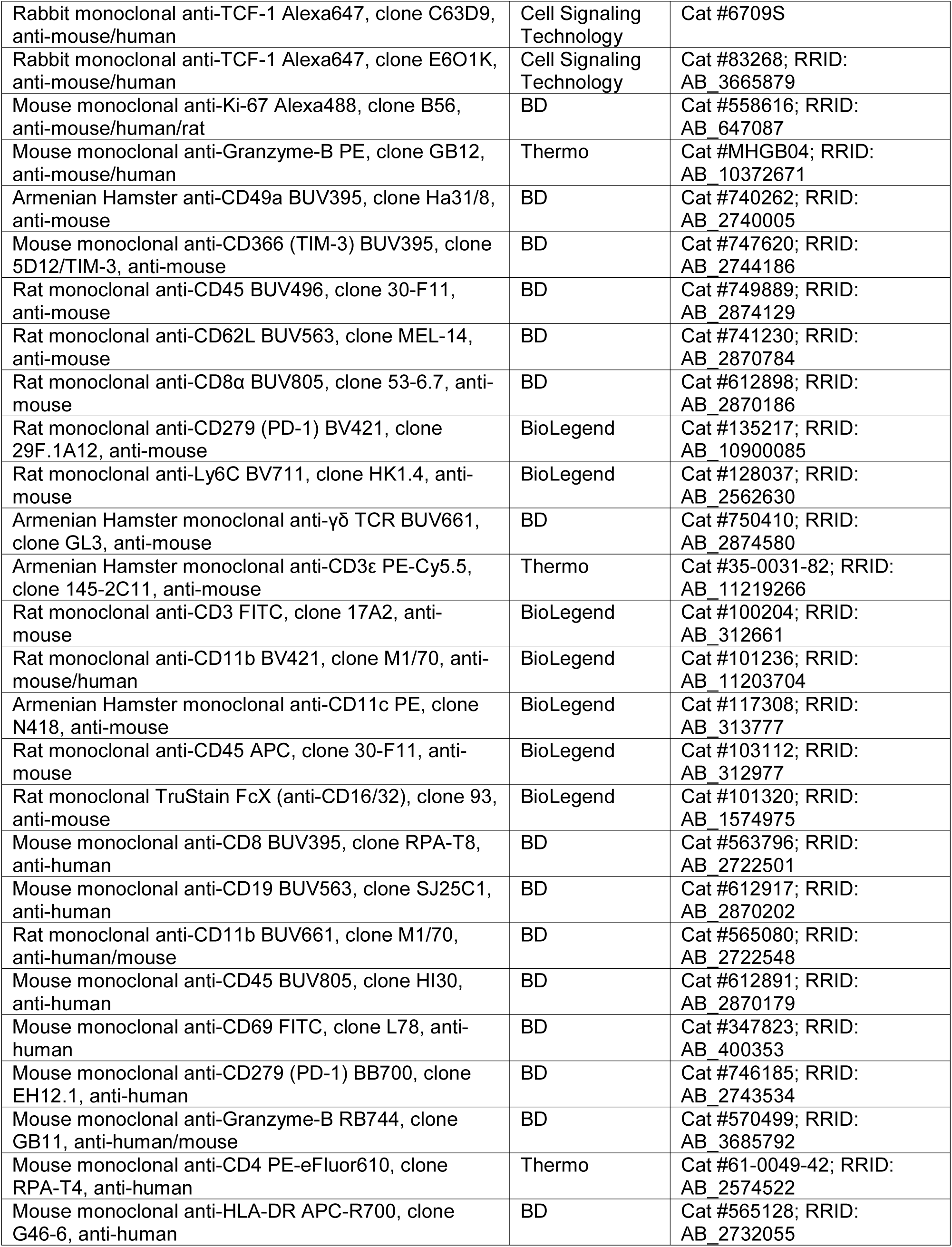
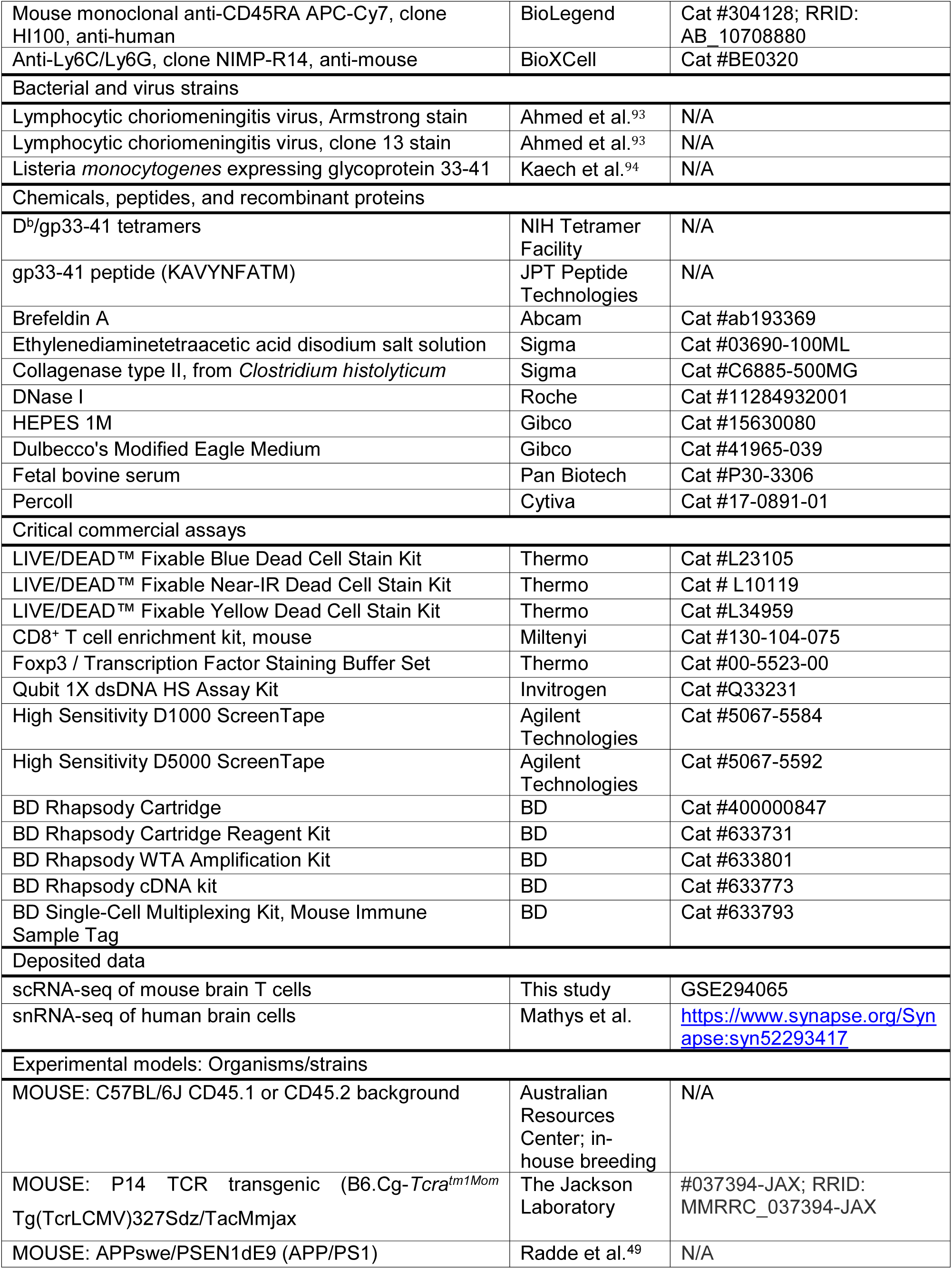
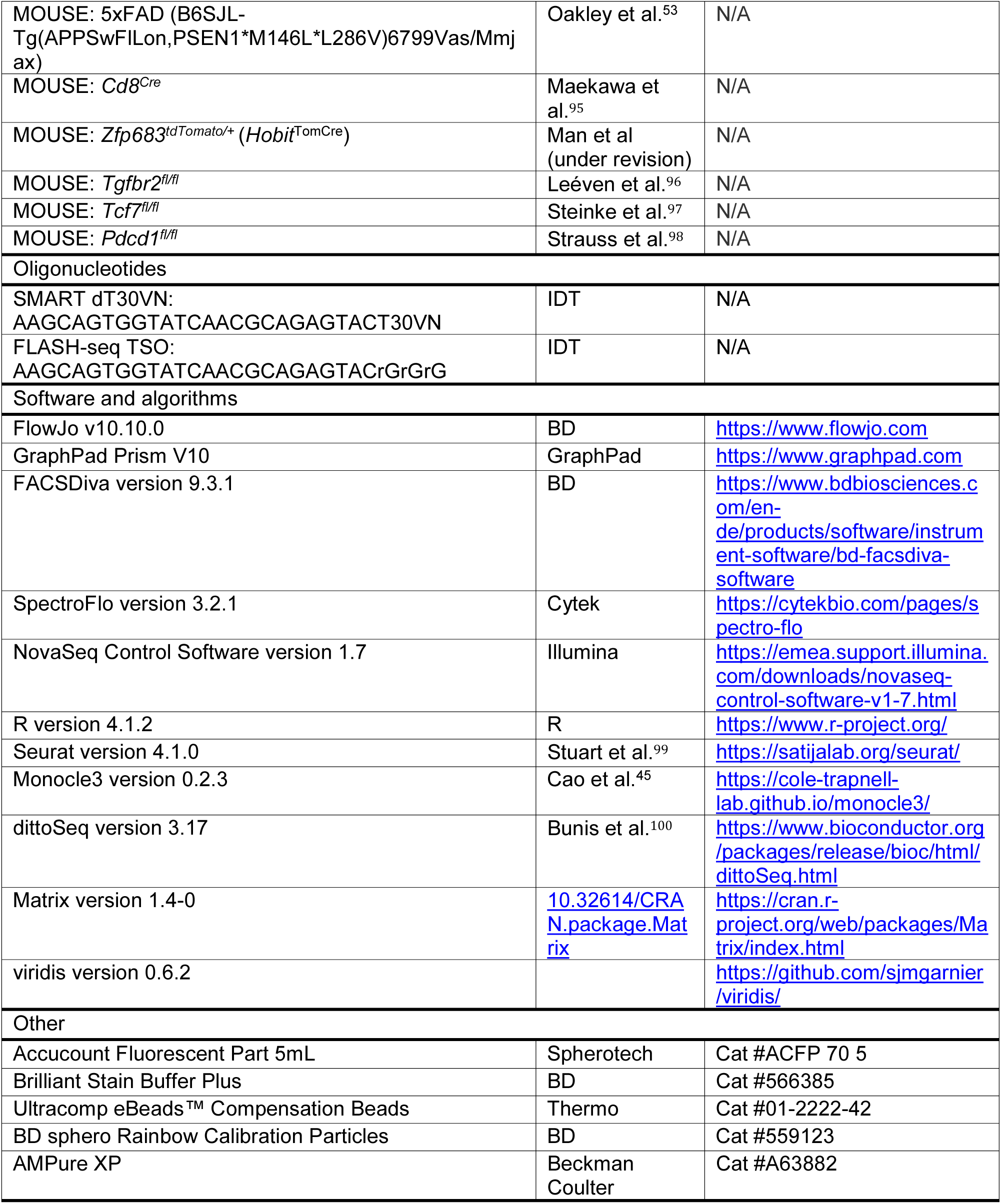

## STAR METHODS

### EXPERIMENTAL MODELS AND STUDY PARTICIPANT DETAILS

#### Mice

Wild-type C57BL/6J mice were bred at the animal facility of the German Center for Neurodegenerative Diseases (Bonn) or at the Peter Doherty Institute for Infection and Immunity (Melbourne). B6.SJL-PtprcaPep3b/BoyJ (CD45.1) and B6.SJL-PtprcaPep3b/BoyJ C57BL/6 (CD45.1/CD45.2) were bred at the Peter Doherty Institute of Infection and Immunity animal facility or obtained from the Australian Resources Centre. APPswe/PSEN1dE9 (APP/PS1) mice^49^ were provided by Martin Fuhrmann. 5xFAD (B6SJL-Tg(APPSwFlLon,PSEN1*M146L*L286V)6799Vas/Mmjax) mice^53^ were provided by Anja Schneider (DZNE Bonn) and backcrossed on C57BL/6J background for 7 generations before use. P14 mice express a transgenic TCR specific for the LCMV immunodominant glycoprotein-derived epitope gp33-41 in the context of H2-D^b^. *E8I^Cre^* (*Cd8^Cre^*) mice express Cre recombinase under joint control of the *Cd8a* promoter and E8I enhancer, rendering Cre expression specific to peripheral CD8^+^ T cells and absent at the CD4^+^ CD8^+^ double-positive thymocyte stage.^95^ *Cd8^Cre^* mice were crossed to *Tgfbr2^fl/fl^* (ref. 96) or *Tcf7^fl/fl^* (ref. 97) mice to induce CD8^+^ T cell-specific deletion of TGF-βRII or TCF-1 (encoded by *Tcf7*), respectively. *Zfp683^tdTomato-Cre/+^* (*Hobit*^TomCre^) mice were generated at the Melbourne Advanced Genome Editing Centre (WEHI) and are described in a separate manuscript. *Tgfbr2^fl/fl^*, *Tcf7^fl/fl^*, and *Pdcd1^fl/fl^* (ref. 98) mice were crossed to *Hobit*^TomCre^ mice. Constitutive *Pdcd1* KO mice were provided by Daniel Gray (WEHI). All mice were maintained on a C57BL/6J background. For experiments involving LCMV infection, mice were used at 10-20 weeks of age; otherwise, mouse age is specified in the respective figure legend. Experimental groups were sex-matched in all experiments performed, and both male and female mice were used. Experiments were approved by the Local Animal Care Commission of North Rhine-Westphalia (licenses 81-02.04.2020.A352 and 81-02.04.2021.A323) or the University of Melbourne Animal Ethics Committee.

#### Human post-mortem brain tissue

Human post-mortem brain tissue samples were obtained from the DZNE Brain Bank with ethic approval from University Hospital Bonn (reference: 229/22). The DZNE Brain Bank obtained permission from the donors for brain autopsy and the use of tissue and clinical information for research purposes. Brain samples (frontal cortex) were isolated postmortem from >70 years-old male patients diagnosed with frontotemporal dementia with tauopathy.

### METHOD DETAILS

#### Human brain sample isolation and processing

Human post-mortem brain tissue samples were directly placed in 9.5 ml of Dulbecco’s Modified Eagle Medium (DMEM; Gibco 41965-039) supplemented with fetal bovine serum (FBS) at 10%. Brain tissue was minced, followed by enzymatic digestion in 10 ml of 10% FBS/DMEM containing collagnease type II (1 mg/ml; Sigma, C6885) and DNase type I (0.12 mg/ml; Roche, 11284932001), in a gentleMACS C tube (Miltenyi, 130-093-237) mounted on a gentleMACS Octo Dissociator with Heaters, selecting the “ABDK_1” program (Miltenyi). Next, EDTA was added to inhibit the enzymatic activity (final concentration 0.5 mM). Brain tissue was further homogenized through a 70 µm strainer using a plunger into a 50 ml conical tube. After centrifugation at 400 g and 4 °C for 5 min, myelin and debris were removed via density centrifugation using a 40% percoll gradient at 1800 g and 18 °C for 20 min without brake. After pelleting, red blood cell lysis was performed using 5 ml of a hypotonic ammonium chloride-potassium bicarbonate (ACK) buffer (in-house), incubated for 1 min at room temperature followed by washing with 20 ml of 1x phosphate buffered saline (PBS; Sigma, D8537-500ML). Cells were then transferred into a 96-well plate (Sarstedt 83.3925500), pelleted, and resuspended in 50 µL of 1x PBS containing fluorescently labelled antibodies targeting surface antigens for 30 min at 4 °C in the dark, in the presence of BD Brilliant Stain Buffer Plus (BD 566385) to minimize physicochemical interactions between fluorochromes. The following anti-human surface antibodies were used: CD8-BUV395 (1:200, BD 563796), CD19-BUV563 (1:200, BD 612917), CD11b-BUV661 (1:200, BD 565080), CD45-BUV805 (1:200, BD 612891), CD69-FITC (1:200, BD 347823), PD-1-BB700 (1:200, BD 746185), CD4-PE-eFluor610 (1:200, Invitrogen 61-0049-42), HLA-DR-APCR700 (1:200, BD 565128), CD45RA-APC-Cy7 (1:200, BioLegend 304128), Fc Blocking Reagent (1:100, Miltenyi 130-059-901), Live/Dead Fixable Yellow Dead Cell Stain (1:1000, Thermo Fischer L34959). Following washing and pelleting, cells were fixed/permeabilized using the eBioscience Transcription Factor Staining kit (eBioscience 00-5523-00) according to the manufacturer instructions. Cells were resuspended in the fixative agent for 1 hr at room temperature, followed by washing twice in 1x perm buffer. This was followed by incubation with antibodies targeting intracellular transcription factors for 2 hr at room temperature. The following antibodies were used at a final volume of 50 μL including 1x perm buffer: Granzyme B-RB744 (1:200, BD 570499), and TCF1-Alexa 647 (1:50, Cell Signaling Technology 83268). Samples were acquired on FACSSymphony S6 instrument (BD) and data was analysed using FlowJo version 10.10.

#### Generation of mixed bone-marrow chimera mice

CD45.1^+^ CD45.2^+^ donor and *Pdcd1* KO CD45.2^+^ donor mice were sacrificed, and bone marrow was flushed from tibia and femur. CD45.1^+^ recipient mice were irradiated (2 doses of 550 Rad), and on the same day were reconstituted with bone marrow from CD45.1^+^ CD45.2^+^ and *Pdcd1* KO CD45.2^+^ mice, mixed at 1:1 ratio, for a total of 4 million cells via the tail vein. Eight weeks later, blood was sampled to confirm reconstitution. Irradiated mice were provided with neomycin supplemented drinking water for the first 4 weeks post-reconstitution.

#### Adoptive T cell transfer

Spleen was isolated from naive P14 donor mice, mashed through a 70µm cell strainer (Miltenyi, cat #130-098-462) into a single-cell suspension, and red blood cells were lysed using ACK buffer (in-house). P14 T cells were enriched from total splenic cells using CD8^+^ T cell enrichment kit (Miltenyi, cat #130-104-075). P14 T cells were adoptively transferred intravenously at a total number of 5x10^4^ cells, namely 2.5x10^4^ per cell population in co-transfer experiments at 1:1 ratio into congenically distinct recipient mice.

#### Viral and bacterial infections

LCMV Armstrong and clone-13 strains were propagated in baby hamster kidney (BHK) cells and titrated on Vero African green monkey kidney cells. BHK cells were inoculated with virus at a multiplicity of infection of 0.05 and incubated at 37°C and 5% CO2. The flask was shaken every 15 min for the first 1.5 hr of incubation, and then incubation was maintained for a total for 48 hr. At 48 hr, the supernatant–containing viral particles–was collected in 500 µL aliquots in cryotubes and frozen at -80 °C until usage. On the day of infection, frozen stocks were diluted in PBS, and mice were injected with 2 x 10^5^ plaque forming units (PFU) intraperitoneally, or with 2 x 10^6^ PFU of LCMV-clone-13 intravenously.

*Listeria monocytogenes* expressing the LCMV immunodominant epitope glycoprotein 33-41 (LM-gp33) was stored in a glycerol stock at -80 °C. A sterile inoculation loop was used to draw a sample from the glycerol stock of LM-gp33, and then to streak a Brain Heart Infusion (BHI) agar plate containing 50 µg/ml streptomycin. Following overnight incubation at 37°C and 5% CO2, a colony of LM-gp33 from the agar plate was picked and then inoculated into 20 ml of BHI broth–containing 50 µg/ml of streptomycin–for ∼18 hr. An aliquot of the bacterial culture was diluted 1:1000 in a final volume of 25 ml BHI broth, and incubated for 3 hr. Optical density was assessed using Spectrophotometer BioMate 3 (Thermo Spectronic) at 600 nm, and incubation was stopped once the OD value exceeded 0.1. Bacterial culture was then serially diluted in PBS, and each mouse received 200 µL via the tail vein, corresponding to 1 x 10^5^ CFU.

For rechallenge experiments with LM-gp33, mice received 200 µg of anti-Gr1 antibody i.p. (BioXCell, BE0320; clone NIMP-R14) 5-7 days before rechallenge (specified in each relevant figure).

#### Labeling of intravascular leukocytes

To label intravascular immune cells, mice received an intravenous injection of 3 μg of CD45-PE-Cy5 or CD45-BV605 (BD, clone 30-F11) antibody in a final volume of 200 μL PBS. Three minutes later, mice were euthanized. For experiments involving PD-1 KO chimeras and P14 co-transfer, a mixture of CD45.1-BV605 (BioLegend, clone A20) and CD45.2-BV605 (BioLegend, clone 104) antibodies (1.5 μg each) was injected.

#### Mouse tissue processing for flow cytometry

Mice were sacrificed via CO_2_ asphyxia or an i.p. injection of ketamine (100 mg/Kg bodyweight) and xylazine (10 mg/Kg bodyweight). The head was cut off; fur, skin, and skull cap were removed, followed by isolation of the brain without inclusion of the dura mater. Brain was minced in a 50 ml tube containing 4 ml of digestion buffer (1 mg/ml collagenase type II, 0.02 mg/ml DNase I, 10 mM HEPES, 9% FBS/DMEM) and incubated for 30 min at 37 °C shaking at 180 rpm. Next, 40 µL of EDTA was added (final concentration of 5 mM) to inhibit the enzymatic activity, together with 10 µL of counting beads. Samples were filtered through a 70-µm strainer into a fresh 50 ml tube, and tissue clumps on the strainer were gently mashed, followed by washing the strainer with 10 ml of 9% FBS/DMEM. Samples were spun down at 300 g 7 min 4 °C, followed by aspiration of supernatant using a suction pump. Cell pellet was resuspended in 10 ml of 27% Percoll, and centrifuged at 600 g for 5 min at 4 °C while setting the brake to 1 (on a scale from 0 to 9). Myelin and debris were aspirated, followed by resuspension in 10 ml of 9% FBS/DMEM. After pelleting and removal of supernatant, the cell pellet was resuspended in 200 µL FACS buffer (3% FBS in 1x PBS) and transferred to a U-bottom 96-well plate for staining with fluorescent antibodies (see “Antibody and tetramer staining of murine T cells” section).

For isolation of lymphocytes from spleen, spleens were mashed over a 70 µm strainer plunged in a 6-well plate filled with 5 ml of PBS. Cell suspension was then transferred to a 50 ml tube, spun down at 400 g 4 °C for 5 min, and supernatant was aspirated using a suction pump. Pellet was resuspended in 1 ml of red blood cell lysis ACK buffer (in-house), incubated at room temperature for 1 min, and the sample was then diluted using 20 ml of PBS. After spinning down and removal of supernatant, cells were resuspended in 2 ml of FACS buffer, and 1/20^th^ of splenic cells were transferred to a U-bottom 96-well plate for staining.

#### Antibody and tetramer staining of murine T cells

Single-cell suspensions were generated as described in the “tissue processing for flow cytometry” section and transferred into a U-bottom 96-well plate (Sarstedt 83.3925500). After pelleting, cells were resuspended in 50 µL anti-mouse Fc block (BioLegend 101319; 1:200) and fixable Live Dead viability near-infrared dye (Thermo, L34975; 1:1000) in PBS for 20 min at 4 °C. Cells were then washed in PBS and incubated with different combinations of fluorescently labelled antibodies targeting surface antigens for 30 min at 4 °C in the presence of BD Brilliant Stain Buffer Plus (BD 566385) to minimize physicochemical interactions between fluorochromes. Different combinations of the following anti-mouse surface antibodies were used in a final volume of 40 µL (including PBS and BD Brilliant Stain Buffer Plus): CD44-BUV395 (1:200, BD 740215), CD4-BUV496 (1:400, BD 612952), TCRβ-BUV563 (1:200, BD 748406), CD49a-BUV661 (1:200, BD 741549), CD69-BUV737 (1:100, BD 564684), CD45.2-BUV737 (1:200, BD 612778), CD62L-BV650 (1:400, BD 564108), PD-1-BV711 (1:200, BioLegend 135231), CD45.1-FITC (1:200, Invitrogen 11-0453-82), KLRG1-BV711 (1:200, BD 564014), CD69-PE-Cy7 (1:200, Invitrogen 25-0691-82), Ly6C-eFluor450 (1:200, Invitrogen 48-5932-82), CX3CR1-BV785 (1:200, BioLegend 149029), CD45.2-BUV805 (1:200, BD 741957), CD8a-PerCP-Cy5.5 (1:100, Invitrogen 45-0081-82), CD103-APC-R700 (1:200, BD 565529), CD8α-BV785 (1:200, BioLegend 100750), TIM3-BV785 (1:200, BioLegend 119725), CD49a-BUV395 (1:200, BD 740262), CD45-BUV496 (1:200, BD 749889), CD62L-BUV563 (1:200, BD 741230), CD8α-BUV805 (1:200, BD 564920), PD-1-BV421 (1:200, BioLegend 135217), Ly6C-BV711 (1:200, BioLegend 128037), γδ TCR-BUV661 (1:200, BD 750410), CD3e-PE-Cy5.5 (1:200, Life Technologies 35-0031-82), CD3-FITC (1:200, BioLegend 100204), CD11b-BV421 (1:200, BioLegend 101236), CD11c-PE (1:200, BioLegend 117308), CD45-APC (1:200, BioLegend 103112).

For experiments involving LCMV infection, CD8^+^ T cells specific for the LCMV immunodominant epitope gp33-41 were characterized using MHC class I tetramers loaded with the gp33-41 peptide. After extracellular staining with antibodies, cells were washed once with PBS, and resuspended in 50 µL of gp33 tetramers diluted 1:400 in FACS buffer. Cells were incubated with the tetramers for 45 min at 4°C, followed by washing in PBS and fixation for subsequent intracellular staining.

For intracellular staining, cells were fixed using the eBioscience Transcription Factor Staining kit (eBioscience 00-5523-00) according to the manufacturer’s instructions. Cells were resuspended in the fixative agent for 1 hr at room temperature, followed by washing twice in 1x perm buffer. This was followed by incubation with antibodies targeting intracellular transcription factors/cytokines for 2 hr at room temperature. For experiments involving *Hobit^Cre^* mice, cells were first fixed in 4% PFA for 20 min at 4 °C to preserve tdTomato’s fluorescence, followed by washing once in PBS, before proceeding to using the eBioscience kit for transcription factor staining as stated above. Different combinations of the following anti-mouse antibodies were used for intracellular staining in a final volume of 30 µL (including 1x perm buffer): Bcl-2 PE (1:100, BD 556537), IFNγ-PE-Cy7 (1:400, Invitrogen 25-7311-82), TNF-APC (1:200, Invitrogen 17-7321-82), TCF1-Alexa 488 (1:100, Cell Signaling Technology 6444S), TCF1-Alexa 647 (1:100, Cell Signaling Technology 6709S), TCF1-Alexa 647 (1:50, Cell Signaling Technology 83268), Ki67-Alexa 488 (1:200, BD 558616), Granzyme B-PE (1:200, Invitrogen MHGB04).

Single-stained UltraComp Compensation Beads were used to set up the compensation matrix (conventional cytometer) or for unmixing (spectral cytometer). Samples were acquired on a FACSSymphony A5 (BD) or Cytek Aurora (Cytek) instrument. For BD FACSSymphony instruments, calibration was performed using CS&T beads for laser delay and 8-peak beads for PMT voltage optimization. For Cytek Aurora, SpectroFlo QC beads were used for calibration.

#### *Ex-vivo* stimulation of murine T cells

Cells were isolated as described under “Mouse tissue processing for flow cytometry” and subjected to surface staining as described under “Antibody and tetramer staining”. For LCMV experiments, cells were restimulated with gp33-41 peptide in T cell medium at a concentration of 5 µM for 30 min at 37 °C and 5% CO_2_. This was followed by the addition of brefeldin A (final dilution 1:1000) and incubation for 4 hr at 37 °C and 5% CO_2_. For PMA/ionomycin stimulation, cells were stimulated for 4 hr at 37 °C and 5% CO_2_ using eBioscience Cell Stimulation Cocktail (500X), and simultaneously adding BD Golgi Plug and Golgi Stop, according to the manufacturer’s instructions. Cells were then washed once using PBS, followed by incubation with Live Dead near IR viability dye (1:1000 in PBS) for 15 min, followed by fixation using the eBioscience kit and intracellular staining as described under “Antibody and tetramer staining of murine T cells”.

#### Cell sorting for scRNA-seq

*BD Rhapsody*: Equivalent amounts (0.05 µg) of a unique hashtag oligo (HTO)-conjugated anti-CD45 antibody (BD Mouse Immune Single-Cell Multiplexing Kit) and fluorescently labelled CD45 antibody were added to each sample during the extracellular staining step described above. Following a 30 min incubation at 4°C, cells were washed, pelleted, and resuspended in FACS buffer in preparation for sorting. Drop delay was set semi-automatically using BD AccuDrop beads. Sorting of extravascular (i.e. negative for the intravenously administered CD45 antibody) live single CD11b^-^ CD45^+^ CD3^+^ was carried out using the 4-way purity sorting mode and 85 µm nozzle of FACSymphony S6 or FACSAria III (BD Bioscience). Cells were collected into BD Sample Buffer in a 1.5 ml tube cooled at 4°C. At the end of the sort, the sample volume was topped up to 615 µL using Sample Buffer as per the manufacturer’s recommendations. Cells were then processed as detailed in the “BD Rhapsody: cell loading, library preparation, and sequencing” section.

Smart-seq2 and FLASH-seq: following extracellular staining and cell washing, cells were resuspended in FACS buffer. An initial round of presorting of non-myeloid cells was carried out using the 4-way purity sort mode, by gating on extravascular CD11b-CD45+ cells. Cells were collected into 9% FBS/DMEM in a 5ml tube. The pre-sorted non-myeloid cells were used to sort CD3^+^ cells using the “Single-Cell” sort mode of FACSAria III or FACSymphony S6 (BD) into 384-well plates containing 1 or 2.3 µL lysis buffer (see below). The plate is then sealed with aluminum foil (VWR 391-1281), spun down at 600 g 4°C for 2 min, and stored at -80°C until processing.

#### BD Rhapsody: cell loading, library preparation, and sequencing

The BD Rhapsody platform is a micro-well-based system that makes use of DNA-barcoded beads, with each oligonucleotide comprising a PCR handle, cell barcode (common to all oligos per bead), unique molecular identifier (unique to each oligo), and oligodT primer at the 3’ end. Cell and bead loading, cell lysis, mRNA-bound bead recovery, reverse transcription and exonuclease treatment were carried out using the BD Rhapsody Express Single-Cell Analysis System according to the manufacturer’s instructions (BD Biosciences). Whole-transcriptome library preparation (based on the random-priming strategy) and index PCR steps were performed using the BD Rhapsody mRNA Whole Transcriptome Analysis and Sample Tag Library Preparation protocol, according to the manufacturer’s recommendations (BD Biosciences). The cDNA libraries were quantified using a Qubit fluorometer with the Qubit double-stranded DNA high-sensitivity kit (Thermo Fisher Scientific), whereas the size distribution of the libraries was assessed using the Agilent high-sensitivity D5000 assay on a TapeStation 4200 system (Agilent Technologies). Paired-end sequencing (2 × 75 cycles) was performed on a NextSeq 500 system (Illumina) using NextSeq 500/550 High Output Kit (version 2.5).

#### Smart-seq2: library preparation and sequencing

Single cells sorted into 384-well plates containing 2.3 µL Smart-seq2 lysis buffer were processed according to the protocol published by Picelli et al^101^ with minor modifications. To ensure reproducibility and pipetting accuracy, a Freedom Evo robot handler (Tecan) was used in all steps, namely reverse transcription, PCR amplification, tagmentation, and index PCR. Lysis buffer contained guanidine hydrochloride (40 mM) instead of Triton-X100 and no RNase inhibitor. During reverse transcription, a template-switch oligo (TSO) is added to allow for the incorporation of a common 5’ barcode at the end of the cDNA molecules. This is followed by PCR amplification using the ISPCR primer, which is complementary to barcodes at both the 5’ and 3’ ends. Full-length cDNA molecules are fragmented and ligated with adapters using the Tn5 transposase enzyme, followed by a final Index PCR step to ligate sequencing adaptors and cell-specific indices. Size distribution of the libraries was assessed using the Agilent High-Sensitivity D5000 assay on a TapeStation 4200. Single-end sequencing was performed on a NovaSeq6000 System using NovaSeq 6000 S1 Reagent Kit.

#### FLASH-seq: library preparation and sequencing

Cells were processed according to the FLASH-seq protocol recently published by Hahaut et al^102^ (https://www.protocols.io/view/flash-seq-protocol-kxygxzkrwv8j/v3). Single cells were sorted into 384-well plates containing 1 µL of lysis buffer per well: Triton-X100 (0.2% v/v; Sigma X100-100ML), dNTP (6 mM each; Carl Roth K039.2), SMART dT30VN (5′Bio-AAGCAGTGGTATCAACGCAGAGTACT30VN-3′ (Bio, biotin), 1.8 µM; IDT), RNase inhibitor (1.2 U/µL; Takara 2313B), DTT (1.2 mM; Thermo 18090050); FLASH-seq TSO (5′Bio-AAGCAGTGGTATCAACGCAGAGTACrGrGrG-3′ (Bio, biotin); 9.2 µM, IDT), dCTP (9 mM; Thermo 10217016), betaine (1 M; Sigma), and nuclease-free water. Plates were stored at -80 °C until further processing. On the day of processing, plate was incubated on a thermocycler at 72 °C for 3 min, followed by addition of 4 µL of reverse transcription-PCR (RT-PCR) mix per well: DTT (4.8 Mm), MgCl_2_ (9.2 mM; Invitrogen AM9530G); betaine (800 mM), RNase inhibitor (0.8 U/µL), SuperScript IV reverse transcriptase (2 U/µL; Thermo 18090050), KAPA HiFi HotStart ReadyMix 2x (1x; Roche KK2602), and nuclease-free water. This mix allows for combining reverse transcription and cDNA PCR-amplification into one reaction^102^ using the following program: 60 min at 50 °C; 98 °C for 3 min, then 21 cycles of (98 °C for 20 s, 67 °C for 20 s, 72 °C for 6 min). cDNA product was purified using AMPure XP beads (Beckman Coulter; A63880) added at a ratio of 0.8x relative to the cDNA product volume. Tagmentation was performed by adding 1.5 µL of loaded Tn5 enzyme (pre-diluted at 1:1250; Diagenode C01070012-30) to 0.5 µL of pre-diluted, normalized cDNA product (∼150 pg/µL). Sample plate was incubated at 55 °C for 8 min followed by stopping the reaction using 1 µL of Nextera XT Neutralize Tagment (NT) buffer (Illumina). For index PCR, 1 µL of a unique dual i5/i7 primer set (IDT) and 3.5 µL of KAPA HiFi HotStart ReadyMix 2x were added per well, followed by applying the following PCR program: 72 °C for 5 min, 98 °C for 30 sec, then 14 cycles of (98 °C for 10 s, 60 °C for 30 s, 72 °C for 30 sec). Single-cell transcriptomes were then pooled from the 384 wells into a single tube, followed by 1x bead purification using AMPure XP beads. Size distribution of the libraries was assessed using the Agilent High-Sensitivity D5000 assay on a TapeStation 4200. Single-end sequencing was performed on a NovaSeq6000 System using NovaSeq 6000 S1 Reagent Kit. An I.DOT instrument (Dispendix) was used for all steps involving addition of small volumes to 384-well plates to ensure accuracy and reproducibility.

#### scRNA-seq data preprocessing and analysis

Raw sequencing files (bcl files) were demultiplexed using the Bcl2fastq2 V2.20 tool from Illumina. Sequencing adapters were trimmed and sequencing reads with a PHRED score >20 were filtered using Cutadapt 1.16. For data generated using BD Rhapsody 3’ assay, STAR aligner^103^ was used to align reads against Gencode vM16 version of mouse reference genome (mm10). Drop-seq tools 2.0.0 were used to generate a unique molecular identifier (UMI)-corrected gene expression count matrix. HTO sequences were added to the reference genome to simultaneously allow for their retrieval during alignment. For data generated using the Smart-seq2 full-length assay, raw reads were pseudo-aligned to the mouse transcriptome using Kallisto^104^ (Gencode vM16 primary assembly) with default settings. Transcripts were quantified as transcript per million reads (TPM) and imported into R using the tximport function of the tximport package (version 1.16.1) while setting the countsFromAbundance argument to “lengthScaledTPM” to account for gene length, given that Smart-seq2 is a full-length scRNA-seq assay.

Downstream analysis was performed in R (version 4.1.2). Rhapsody datasets were filtered using the barcodeRanks() function to exclude cells with UMI counts below the inflection point, which represents the sharp transition in UMI counts between cell-containing wells and empty wells.^105^ Downstream analysis was performed using the Seurat R package (version 4.1.0).^99^ For Rhapsody datasets, the data was further filtered to exclude cells expressing less than 200 genes, more than 2000 genes or cells whose mitochondrial reads account for more than 10% of their transcriptomes. For Smart-seq2 datasets, the data was filtered to exclude cells expressing less than 500 genes, more than 4000 genes or cells whose mitochondrial reads account for more than 10% of their transcriptomes.

Normalization and scaling were performed using the SCTransform() function.^106^ Principal component analysis (PCA) was applied using the RunPCA() function of Seurat, and– based on elbow plot and the inspection of the individual PCs and their contribution to the variance in the data–different number of PCs across datasets were used to run FindNeighbors() funciton. Clustering and non-linear dimensionality reductions were performed using the FindClusters() and RunUMAP() functions, respectively. Differential expression analysis was performed using the FindAllMarkers() function, setting both min.pct and logfc.threshold to 0.2 and using the default Wilcoxon Rank Sum test. Pairwise DEG analysis between conditions across clusters was performed using Seurat’s FindMarkers function in a for-loop, setting min.pct to 0.2 and logfc.threshold to 0.3. Heatmaps were generated by a) computing the average gene expression per cluster using Seurat’s AverageExpression function, setting the slot to “scale.data”, and b) convert AverageExpression output into a matrix datatype and pass it to pheatmap function, setting the *scale* argument to “row”. The “viridis” package was used to set the scale colors in FeaturePlots. To compute TGF-β signaling score, a gene list corresponding to the signaling pathway was imported from a study by Nath et al,^70^ converted into a list datatype, and then Seurat’s AddModuleScore function was used to compute a gene-expression score. Trajectory inference was conducted using the monocle3 package (version 0.2.3).^45^ The cluster_cells function from monocle3 was used while setting k nearest neighbor to 35 and using “Louvain” as the clustering method. Histograms were generated using the dittoSeq package.^100^

#### Statistics

All statistical analyses, except for scRNA-seq data analysis, were performed using GraphPad Prism 9.5.1. For comparisons between two groups across one variable, unpaired two-tailed Student’s *t* test was used. For comparisons between paired data (P14 co-transfer experiments and PD-1 KO chimeras), paired Student’s *t* test was used. For comparison between two groups across two or more variables, two-way Analysis of Variance (ANOVA) with Šídák’s or Tukey’s multiple comparisons test were used. The exact statistical test used for each figure panel is specified in the respective figure legend. Statistical differences were considered significant if α was < 0.05. Data is reported as mean ± standard error of the mean (sem).

## References

1. Zimmermann, C., Prevost-Blondel, A., Blaser, C., and Pircher, H. (1999). Kinetics of the response of naive and memory CD8 T cells to antigen: similarities and differences. Eur J Immunol 29, 284–290. 10.1002/(SICI)1521-4141(199901)29:01<284::AID-IMMU284>3.0.CO;2-C.

2. Cho, B.K., Wang, C., Sugawa, S., Eisen, H.N., and Chen, J. (1999). Functional differences between memory and naive CD8 T cells. Proc Natl Acad Sci U S A 96, 2976–2981. 10.1073/pnas.96.6.2976.

3. Masopust, D., Vezys, V., Marzo, A.L., and Lefrancois, L. (2001). Preferential localization of effector memory cells in nonlymphoid tissue. Science 291, 2413–2417. 10.1126/science.1058867.

4. Gebhardt, T., Wakim, L.M., Eidsmo, L., Reading, P.C., Heath, W.R., and Carbone, F.R. (2009). Memory T cells in nonlymphoid tissue that provide enhanced local immunity during infection with herpes simplex virus. Nat Immunol 10, 524–530. 10.1038/ni.1718.

5. Sallusto, F., Lenig, D., Forster, R., Lipp, M., and Lanzavecchia, A. (1999). Two subsets of memory T lymphocytes with distinct homing potentials and effector functions. Nature 401, 708–712. 10.1038/44385.

6. Gerlach, C., Moseman, E.A., Loughhead, S.M., Alvarez, D., Zwijnenburg, A.J., Waanders, L., Garg, R., de la Torre, J.C., and von Andrian, U.H. (2016). The Chemokine Receptor CX3CR1 Defines Three Antigen-Experienced CD8 T Cell Subsets with Distinct Roles in Immune Surveillance and Homeostasis. Immunity 45, 1270–1284. 10.1016/j.immuni.2016.10.018.

7. Bottcher, J.P., Beyer, M., Meissner, F., Abdullah, Z., Sander, J., Hochst, B., Eickhoff, S., Rieckmann, J.C., Russo, C., Bauer, T., et al. (2015). Functional classification of memory CD8(+) T cells by CX3CR1 expression. Nat Commun 6, 8306 10.1038/ncomms9306.

8. Kaech, S.M., and Cui, W. (2012). Transcriptional control of effector and memory CD8+ T cell differentiation. Nat Rev Immunol 12, 749–761 10.1038/nri3307.

9. Gebhardt, T., Palendira, U., Tscharke, D.C., and Bedoui, S. (2018). Tissue-resident memory T cells in tissue homeostasis, persistent infection, and cancer surveillance. Immunol Rev 283, 54–76. 10.1111/imr.12650.

10. Schenkel, J.M., Fraser, K.A., Beura, L.K., Pauken, K.E., Vezys, V., and Masopust, D. (2014). T cell memory. Resident memory CD8 T cells trigger protective innate and adaptive immune responses. Science 346, 98–101. 10.1126/science.1254536.

11. Schenkel, J.M., Fraser, K.A., Vezys, V., and Masopust, D. (2013). Sensing and alarm function of resident memory CD8(+) T cells. Nat Immunol 14, 509–513. 10.1038/ni.2568.

12. Ariotti, S., Hogenbirk, M.A., Dijkgraaf, F.E., Visser, L.L., Hoekstra, M.E., Song, J.Y., Jacobs, H., Haanen, J.B., and Schumacher, T.N. (2014). T cell memory. Skin-resident memory CD8(+) T cells trigger a state of tissue-wide pathogen alert. Science 346, 101–105. 10.1126/science.1254803.

13. Mackay, L.K., Wynne-Jones, E., Freestone, D., Pellicci, D.G., Mielke, L.A., Newman, D.M., Braun, A., Masson, F., Kallies, A., Belz, G.T., and Carbone, F.R. (2015). T-box Transcription Factors Combine with the Cytokines TGF-beta and IL-15 to Control Tissue-Resident Memory T Cell Fate. Immunity 43, 1101–1111. 10.1016/j.immuni.2015.11.008.

14. Masopust, D., Choo, D., Vezys, V., Wherry, E.J., Duraiswamy, J., Akondy, R., Wang, J., Casey, K.A., Barber, D.L., Kawamura, K.S., et al. (2010). Dynamic T cell migration program provides resident memory within intestinal epithelium. J Exp Med 207, 553–564. 10.1084/jem.20090858.

15. Milner, J.J., Toma, C., Yu, B., Zhang, K., Omilusik, K., Phan, A.T., Wang, D., Getzler, A.J., Nguyen, T., Crotty, S., et al. (2017). Runx3 programs CD8(+) T cell residency in non-lymphoid tissues and tumours. Nature 552, 253–257. 10.1038/nature24993.

16. Milner, J.J., Toma, C., He, Z., Kurd, N.S., Nguyen, Q.P., McDonald, B., Quezada, L., Widjaja, C.E., Witherden, D.A., Crowl, J.T., et al. (2020). Heterogenous Populations of Tissue-Resident CD8(+) T Cells Are Generated in Response to Infection and Malignancy. Immunity 52, 808–824 e807. 10.1016/j.immuni.2020.04.007.

17. Mackay, L.K., Rahimpour, A., Ma, J.Z., Collins, N., Stock, A.T., Hafon, M.L., Vega-Ramos, J., Lauzurica, P., Mueller, S.N., Stefanovic, T., et al. (2013). The developmental pathway for CD103(+)CD8+ tissue-resident memory T cells of skin. Nat Immunol 14, 1294–1301. 10.1038/ni.2744.

18. Topham, D.J., and Reilly, E.C. (2018). Tissue-Resident Memory CD8(+) T Cells: From Phenotype to Function. Front Immunol 9, 515. 10.3389/fimmu.2018.00515.

19. Mackay, L.K., Minnich, M., Kragten, N.A., Liao, Y., Nota, B., Seillet, C., Zaid, A., Man, K., Preston, S., Freestone, D., et al. (2016). Hobit and Blimp1 instruct a universal transcriptional program of tissue residency in lymphocytes. Science 352, 459–463. 10.1126/science.aad2035.

20. Christo, S.N., Evrard, M., Park, S.L., Gandolfo, L.C., Burn, T.N., Fonseca, R., Newman, D.M., Alexandre, Y.O., Collins, N., Zamudio, N.M., et al. (2021). Discrete tissue microenvironments instruct diversity in resident memory T cell function and plasticity. Nat Immunol 22, 1140–1151. 10.1038/s41590-021-01004-1.

21. Crowl, J.T., Heeg, M., Ferry, A., Milner, J.J., Omilusik, K.D., Toma, C., He, Z., Chang, J.T., and Goldrath, A.W. (2022). Tissue-resident memory CD8(+) T cells possess unique transcriptional, epigenetic and functional adaptations to different tissue environments. Nature immunology 23, 1121–1131. 10.1038/s41590-022-01229-8.

22. Hirai, T., Yang, Y., Zenke, Y., Li, H., Chaudhri, V.K., De La Cruz Diaz, J.S., Zhou, P.Y., Nguyen, B.A., Bartholin, L., Workman, C.J., et al. (2021). Competition for Active TGFbeta Cytokine Allows for Selective Retention of Antigen-Specific Tissue-Resident Memory T Cells in the Epidermal Niche. Immunity 54, 84–98 e85. 10.1016/j.immuni.2020.10.022.

23. Casey, K.A., Fraser, K.A., Schenkel, J.M., Moran, A., Abt, M.C., Beura, L.K., Lucas, P.J., Artis, D., Wherry, E.J., Hogquist, K., et al. (2012). Antigen-independent differentiation and maintenance of effector-like resident memory T cells in tissues. J Immunol 188, 4866–4875. 10.4049/jimmunol.1200402.

24. Zhang, N., and Bevan, M.J. (2013). Transforming growth factor-beta signaling controls the formation and maintenance of gut-resident memory T cells by regulating migration and retention. Immunity 39, 687–696. 10.1016/j.immuni.2013.08.019.

25. Thom, J.T., Weber, T.C., Walton, S.M., Torti, N., and Oxenius, A. (2015). The Salivary Gland Acts as a Sink for Tissue-Resident Memory CD8(+) T Cells, Facilitating Protection from Local Cytomegalovirus Infection. Cell Rep 13, 1125–1136. 10.1016/j.celrep.2015.09.082.

26. Bromley, S.K., Akbaba, H., Mani, V., Mora-Buch, R., Chasse, A.Y., Sama, A., and Luster, A.D. (2020). CD49a Regulates Cutaneous Resident Memory CD8(+) T Cell Persistence and Response. Cell Rep 32, 108085. 10.1016/j.celrep.2020.108085.

27. Wakim, L.M., Woodward-Davis, A., and Bevan, M.J. (2010). Memory T cells persisting within the brain after local infection show functional adaptations to their tissue of residence. Proc Natl Acad Sci U S A 107, 17872–17879. 10.1073/pnas.1010201107.

28. Wakim, L.M., Woodward-Davis, A., Liu, R., Hu, Y., Villadangos, J., Smyth, G., and Bevan, M.J. (2012). The molecular signature of tissue resident memory CD8 T cells isolated from the brain. J Immunol 189, 3462–3471. 10.4049/jimmunol.1201305.

29. Steinbach, K., Vincenti, I., Kreutzfeldt, M., Page, N., Muschaweckh, A., Wagner, I., Drexler, I., Pinschewer, D., Korn, T., and Merkler, D. (2016). Brain-resident memory T cells represent an autonomous cytotoxic barrier to viral infection. J Exp Med 213, 1571–1587. 10.1084/jem.20151916.

30. Wherry, E.J., Blattman, J.N., Murali-Krishna, K., van der Most, R., and Ahmed, R. (2003). Viral persistence alters CD8 T-cell immunodominance and tissue distribution and results in distinct stages of functional impairment. J Virol 77, 4911–4927. 10.1128/jvi.77.8.4911-4927.2003.

31. Urban, S.L., Jensen, I.J., Shan, Q., Pewe, L.L., Xue, H.H., Badovinac, V.P., and Harty, J.T. (2020). Peripherally induced brain tissue-resident memory CD8(+) T cells mediate protection against CNS infection. Nature immunology 21, 938–949. 10.1038/s41590-020-0711-8.

32. Maru, S., Jin, G., Schell, T.D., and Lukacher, A.E. (2017). TCR stimulation strength is inversely associated with establishment of functional brain-resident memory CD8 T cells during persistent viral infection. PLoS Pathog 13, e1006318. 10.1371/journal.ppat.1006318.

33. Groh, J., Knöpper, K., Arampatzi, P., Yuan, X., Lößlein, L., Saliba, A.-E., Kastenmüller, W., and Martini, R. (2021). Accumulation of cytotoxic T cells in the aged CNS leads to axon degeneration and contributes to cognitive and motor decline. Nat Ageing, 357-367.

34. Su, W., Saravia, J., Risch, I., Rankin, S., Guy, C., Chapman, N.M., Shi, H., Sun, Y., Kc, A., Li, W., et al. (2023). CXCR6 orchestrates brain CD8(+) T cell residency and limits mouse Alzheimer’s disease pathology. Nat Immunol 24, 1735–1747. 10.1038/s41590-023-01604-z.

35. Chen, X., Firulyova, M., Manis, M., Herz, J., Smirnov, I., Aladyeva, E., Wang, C., Bao, X., Finn, M.B., Hu, H., et al. (2023). Microglia-mediated T cell infiltration drives neurodegeneration in tauopathy. Nature 615, 668–677. 10.1038/s41586-023-05788-0.

36. Vincenti, I., Page, N., Steinbach, K., Yermanos, A., Lemeille, S., Nunez, N., Kreutzfeldt, M., Klimek, B., Di Liberto, G., Egervari, K., et al. (2022). Tissue-resident memory CD8(+) T cells cooperate with CD4(+) T cells to drive compartmentalized immunopathology in the CNS. Sci Transl Med 14, eabl6058. 10.1126/scitranslmed.abl6058.

37. Ayasoufi, K., Wolf, D.M., Namen, S.L., Jin, F., Tritz, Z.P., Pfaller, C.K., Zheng, J., Goddery, E.N., Fain, C.E., Gulbicki, L.R., et al. (2023). Brain resident memory T cells rapidly expand and initiate neuroinflammatory responses following CNS viral infection. Brain Behav Immun 112, 51–76. 10.1016/j.bbi.2023.05.009.

38. Pasciuto, E., Burton, O.T., Roca, C.P., Lagou, V., Rajan, W.D., Theys, T., Mancuso, R., Tito, R.Y., Kouser, L., Callaerts-Vegh, Z., et al. (2020). Microglia Require CD4 T Cells to Complete the Fetal-to-Adult Transition. Cell 182, 625–640 e624. 10.1016/j.cell.2020.06.026.

39. Smolders, J., Heutinck, K.M., Fransen, N.L., Remmerswaal, E.B.M., Hombrink, P., Ten Berge, I.J.M., van Lier, R.A.W., Huitinga, I., and Hamann, J. (2018). Tissue-resident memory T cells populate the human brain. Nat Commun 9, 4593. 10.1038/s41467-018-07053-9.

40. Mrdjen, D., Pavlovic, A., Hartmann, F.J., Schreiner, B., Utz, S.G., Leung, B.P., Lelios, I., Heppner, F.L., Kipnis, J., Merkler, D., et al. (2018). High-Dimensional Single-Cell Mapping of Central Nervous System Immune Cells Reveals Distinct Myeloid Subsets in Health, Aging, and Disease. Immunity 48, 599. 10.1016/j.immuni.2018.02.014.

41. Anderson, K.G., Mayer-Barber, K., Sung, H., Beura, L., James, B.R., Taylor, J.J., Qunaj, L., Griffith, T.S., Vezys, V., Barber, D.L., and Masopust, D. (2014). Intravascular staining for discrimination of vascular and tissue leukocytes. Nat Protoc 9, 209–222. 10.1038/nprot.2014.005.

42. Zhou, X., Yu, S., Zhao, D.M., Harty, J.T., Badovinac, V.P., and Xue, H.H. (2010). Differentiation and persistence of memory CD8(+) T cells depend on T cell factor 1. Immunity 33, 229–240. 10.1016/j.immuni.2010.08.002.

43. Jeannet, G., Boudousquie, C., Gardiol, N., Kang, J., Huelsken, J., and Held, W. (2010). Essential role of the Wnt pathway effector Tcf-1 for the establishment of functional CD8 T cell memory. Proc Natl Acad Sci U S A 107, 9777–9782. 10.1073/pnas.0914127107.

44. Pais Ferreira, D., Silva, J.G., Wyss, T., Fuertes Marraco, S.A., Scarpellino, L., Charmoy, M., Maas, R., Siddiqui, I., Tang, L., Joyce, J.A., et al. (2020). Central memory CD8(+) T cells derive from stem-like Tcf7(hi) effector cells in the absence of cytotoxic differentiation. Immunity 53, 985–1000 e1011. 10.1016/j.immuni.2020.09.005.

45. Cao, J., Spielmann, M., Qiu, X., Huang, X., Ibrahim, D.M., Hill, A.J., Zhang, F., Mundlos, S., Christiansen, L., Steemers, F.J., et al. (2019). The single-cell transcriptional landscape of mammalian organogenesis. Nature 566, 496–502. 10.1038/s41586-019-0969-x.

46. Mogilenko, D.A., Shpynov, O., Andhey, P.S., Arthur, L., Swain, A., Esaulova, E., Brioschi, S., Shchukina, I., Kerndl, M., Bambouskova, M., et al. (2021). Comprehensive Profiling of an Aging Immune System Reveals Clonal GZMK(+) CD8(+) T Cells as Conserved Hallmark of Inflammaging. Immunity 54, 99–115 e112. 10.1016/j.immuni.2020.11.005.

47. Krishnarajah, S., Ingelfinger, F., Friebel, E., Cansever, D., Amorim, A., Andreadou, M., Bamert, D., Litscher, G., Lutz, M., Mayoux, M., et al. (2022). Single-cell profiling of immune system alterations in lymphoid, barrier and solid tissues in aged mice. Nat Aging 2, 74–89. 10.1038/s43587-021-00148-x.

48. Zhang, S., Gao, Y., Zhao, Y., Huang, T.Y., Zheng, Q., and Wang, X. (2025). Peripheral and central neuroimmune mechanisms in Alzheimer’s disease pathogenesis. Mol Neurodegener 20, 22. 10.1186/s13024-025-00812-5.

49. Radde, R., Bolmont, T., Kaeser, S.A., Coomaraswamy, J., Lindau, D., Stoltze, L., Calhoun, M.E., Jaggi, F., Wolburg, H., Gengler, S., et al. (2006). Abeta42-driven cerebral amyloidosis in transgenic mice reveals early and robust pathology. EMBO Rep 7, 940–946. 10.1038/sj.embor.7400784.

50. Keren-Shaul, H., Spinrad, A., Weiner, A., Matcovitch-Natan, O., Dvir-Szternfeld, R., Ulland, T.K., David, E., Baruch, K., Lara-Astaiso, D., Toth, B., et al. (2017). A Unique Microglia Type Associated with Restricting Development of Alzheimer’s Disease. Cell 169, 1276–1290 e1217. 10.1016/j.cell.2017.05.018.

51. Kamphuis, W., Kooijman, L., Schetters, S., Orre, M., and Hol, E.M. (2016). Transcriptional profiling of CD11c-positive microglia accumulating around amyloid plaques in a mouse model for Alzheimer’s disease. Biochim Biophys Acta 1862, 1847–1860. 10.1016/j.bbadis.2016.07.007.

52. Unger, M.S., Li, E., Scharnagl, L., Poupardin, R., Altendorfer, B., Mrowetz, H., Hutter-Paier, B., Weiger, T.M., Heneka, M.T., Attems, J., and Aigner, L. (2020). CD8(+) T-cells infiltrate Alzheimer’s disease brains and regulate neuronal- and synapse-related gene expression in APP-PS1 transgenic mice. Brain Behav Immun 89, 67–86. 10.1016/j.bbi.2020.05.070.

53. Oakley, H., Cole, S.L., Logan, S., Maus, E., Shao, P., Craft, J., Guillozet-Bongaarts, A., Ohno, M., Disterhoft, J., Van Eldik, L., et al. (2006). Intraneuronal beta-amyloid aggregates, neurodegeneration, and neuron loss in transgenic mice with five familial Alzheimer’s disease mutations: potential factors in amyloid plaque formation. J Neurosci 26, 10129–10140. 10.1523/JNEUROSCI.1202-06.2006.

54. Mathys, H., Peng, Z., Boix, C.A., Victor, M.B., Leary, N., Babu, S., Abdelhady, G., Jiang, X., Ng, A.P., Ghafari, K., et al. (2023). Single-cell atlas reveals correlates of high cognitive function, dementia, and resilience to Alzheimer’s disease pathology. Cell 186, 4365–4385 e4327. 10.1016/j.cell.2023.08.039.

55. Prasad, S., Hu, S., Sheng, W.S., Chauhan, P., Singh, A., and Lokensgard, J.R. (2017). The PD-1: PD-L1 pathway promotes development of brain-resident memory T cells following acute viral encephalitis. J Neuroinflammation 14, 82. 10.1186/s12974-017-0860-3.

56. Jubel, J.M., Barbati, Z.R., Burger, C., Wirtz, D.C., and Schildberg, F.A. (2020). The Role of PD-1 in Acute and Chronic Infection. Front Immunol 11, 487. 10.3389/fimmu.2020.00487.

57. Ahmed, R., Simon, R.S., Matloubian, M., Kolhekar, S.R., Southern, P.J., and Freedman, D.M. (1988). Genetic analysis of in vivo-selected viral variants causing chronic infection: importance of mutation in the L RNA segment of lymphocytic choriomeningitis virus. J Virol 62, 3301–3308. 10.1128/JVI.62.9.3301-3308.1988.

58. Bergthaler, A., Flatz, L., Hegazy, A.N., Johnson, S., Horvath, E., Lohning, M., and Pinschewer, D.D. (2010). Viral replicative capacity is the primary determinant of lymphocytic choriomeningitis virus persistence and immunosuppression. Proc Natl Acad Sci U S A 107, 21641–21646. 10.1073/pnas.1011998107.

59. Utzschneider, D.T., Gabriel, S.S., Chisanga, D., Gloury, R., Gubser, P.M., Vasanthakumar, A., Shi, W., and Kallies, A. (2020). Early precursor T cells establish and propagate T cell exhaustion in chronic infection. Nat Immunol 21, 1256–1266. 10.1038/s41590-020-0760-z.

60. Wu, J., Madi, A., Mieg, A., Hotz-Wagenblatt, A., Weisshaar, N., Ma, S., Mohr, K., Schlimbach, T., Hering, M., Borgers, H., and Cui, G. (2020). T Cell Factor 1 Suppresses CD103+ Lung Tissue-Resident Memory T Cell Development. Cell Rep 31, 107484. 10.1016/j.celrep.2020.03.048.

61. Matsuzaki, J., Tsuji, T., Chamoto, K., Takeshima, T., Sendo, F., and Nishimura, T. (2003). Successful elimination of memory-type CD8+ T cell subsets by the administration of anti-Gr-1 monoclonal antibody in vivo. Cell Immunol 224, 98–105. 10.1016/j.cellimm.2003.08.009.

62. Evrard, M., Becht, E., Fonseca, R., Obers, A., Park, S.L., Ghabdan-Zanluqui, N., Schroeder, J., Christo, S.N., Schienstock, D., Lai, J., et al. (2023). Single-cell protein expression profiling resolves circulating and resident memory T cell diversity across tissues and infection contexts. Immunity 56, 1664–1680 e1669. 10.1016/j.immuni.2023.06.005.

63. Cassidy, B.R., Zhang, M., Sonntag, W.E., and Drevets, D.A. (2020). Neuroinvasive Listeria monocytogenes infection triggers accumulation of brain CD8(+) tissue-resident memory T cells in a miR-155-dependent fashion. J Neuroinflammation 17, 259. 10.1186/s12974-020-01929-8.

64. Ghosh, P., and Higgins, D.E. (2018). Listeria monocytogenes Infection of the Brain. J Vis Exp. 10.3791/58723.

65. Scholler, A.S., Nazerai, L., Christensen, J.P., and Thomsen, A.R. (2020). Functionally Competent, PD-1(+) CD8(+) Trm Cells Populate the Brain Following Local Antigen Encounter. Front Immunol 11, 595707. 10.3389/fimmu.2020.595707.

66. Phares, T.W., Ramakrishna, C., Parra, G.I., Epstein, A., Chen, L., Atkinson, R., Stohlman, S.A., and Bergmann, C.C. (2009). Target-dependent B7-H1 regulation contributes to clearance of central nervous system infection and dampens morbidity. J Immunol 182, 5430–5438. 10.4049/jimmunol.0803557.

67. Shwetank Frost, E.L., Mockus, T.E., Ren, H.M., Toprak, M., Lauver, M.D., Netherby-Winslow, C.S., Jin, G., Cosby, J.M., Evavold, B.D., and Lukacher, A.E. (2019). PD-1 Dynamically Regulates Inflammation and Development of Brain-Resident Memory CD8 T Cells During Persistent Viral Encephalitis. Front Immunol 10, 783. 10.3389/fimmu.2019.00783.

68. Prasad, S., Hu, S., Sheng, W.S., Singh, A., and Lokensgard, J.R. (2015). Tregs Modulate Lymphocyte Proliferation, Activation, and Resident-Memory T-Cell Accumulation within the Brain during MCMV Infection. PLoS One 10, e0145457. 10.1371/journal.pone.0145457.

69. Graham, J.B., Da Costa, A., and Lund, J.M. (2014). Regulatory T cells shape the resident memory T cell response to virus infection in the tissues. J Immunol 192, 683–690. 10.4049/jimmunol.1202153.

80. Liao, W., Liu, Y., Ma, C., Wang, L., Li, G., Mishra, S., Srinivasan, S., Fan, K.K., Wu, H., Li, Q., et al. (2021). The downregulation of IL-18R defines bona fide kidney-resident CD8(+) T cells. iScience 24, 101975. 10.1016/j.isci.2020.101975.

81. Page, N., Lemeille, S., Vincenti, I., Klimek, B., Mariotte, A., Wagner, I., Di Liberto, G., Kaye, J., and Merkler, D. (2021). Persistence of self-reactive CD8+ T cells in the CNS requires TOX-dependent chromatin remodeling. Nat Commun 12, 1009. 10.1038/s41467-021-21109-3.

82. Page, N., Klimek, B., De Roo, M., Steinbach, K., Soldati, H., Lemeille, S., Wagner, I., Kreutzfeldt, M., Di Liberto, G., Vincenti, I., et al. (2018). Expression of the DNA-Binding Factor TOX Promotes the Encephalitogenic Potential of Microbe-Induced Autoreactive CD8(+) T Cells. Immunity 48, 937–950 e938. 10.1016/j.immuni.2018.04.005.

83. Kumar, B.V., Ma, W., Miron, M., Granot, T., Guyer, R.S., Carpenter, D.J., Senda, T., Sun, X., Ho, S.H., Lerner, H., et al. (2017). Human Tissue-Resident Memory T Cells Are Defined by Core Transcriptional and Functional Signatures in Lymphoid and Mucosal Sites. Cell Rep 20, 2921–2934. 10.1016/j.celrep.2017.08.078.

84. Smith, C.J., and Snyder, C.M. (2021). Inhibitory Molecules PD-1, CD73 and CD39 Are Expressed by CD8(+) T Cells in a Tissue-Dependent Manner and Can Inhibit T Cell Responses to Stimulation. Front Immunol 12, 704862. 10.3389/fimmu.2021.704862.

85. Pauken, K.E., Godec, J., Odorizzi, P.M., Brown, K.E., Yates, K.B., Ngiow, S.F., Burke, K.P., Maleri, S., Grande, S.M., Francisco, L.M., et al. (2020). The PD-1 Pathway Regulates Development and Function of Memory CD8(+) T Cells following Respiratory Viral Infection. Cell Rep 31, 107827. 10.1016/j.celrep.2020.107827.

86. Le Moine, M., Azouz, A., Sanchez Sanchez, G., Dejolier, S., Nguyen, M., Thomas, S., Shala, V., Dreidi, H., Denanglaire, S., Libert, F., et al. (2023). Homeostatic PD-1 signaling restrains EOMES-dependent oligoclonal expansion of liver-resident CD8 T cells. Cell Rep 42, 112876. 10.1016/j.celrep.2023.112876.

87. Charlton, J.J., Chatzidakis, I., Tsoukatou, D., Boumpas, D.T., Garinis, G.A., and Mamalaki, C. (2013). Programmed death-1 shapes memory phenotype CD8 T cell subsets in a cell-intrinsic manner. J Immunol 190, 6104–6114. 10.4049/jimmunol.1201617.

88. Wang, Z., Wang, S., Goplen, N.P., Li, C., Cheon, I.S., Dai, Q., Huang, S., Shan, J., Ma, C., Ye, Z., et al. (2019). PD-1(hi) CD8(+) resident memory T cells balance immunity and fibrotic sequelae. Sci Immunol 4. 10.1126/sciimmunol.aaw1217.

89. Spittau, B., Dokalis, N., and Prinz, M. (2020). The Role of TGFbeta Signaling in Microglia Maturation and Activation. Trends Immunol 41, 836–848. 10.1016/j.it.2020.07.003.

90. Ong, C.H., Tham, C.L., Harith, H.H., Firdaus, N., and Israf, D.A. (2021). TGF-beta-induced fibrosis: A review on the underlying mechanism and potential therapeutic strategies. Eur J Pharmacol 911, 174510. 10.1016/j.ejphar.2021.174510.

91. Kaya, T., Mattugini, N., Liu, L., Ji, H., Cantuti-Castelvetri, L., Wu, J., Schifferer, M., Groh, J., Martini, R., Besson-Girard, S., et al. (2022). CD8(+) T cells induce interferon-responsive oligodendrocytes and microglia in white matter aging. Nat Neurosci 25, 1446–1457. 10.1038/s41593-022-01183-6.

92. Jaakkola, I., Merinen, M., Jalkanen, S., and Hanninen, A. (2003). Ly6C induces clustering of LFA-1 (CD11a/CD18) and is involved in subtype-specific adhesion of CD8 T cells. J Immunol 170, 1283–1290. 10.4049/jimmunol.170.3.1283.

93. Ahmed, R., Salmi, A., Butler, L.D., Chiller, J.M., and Oldstone, M.B. (1984). Selection of genetic variants of lymphocytic choriomeningitis virus in spleens of persistently infected mice. Role in suppression of cytotoxic T lymphocyte response and viral persistence. J Exp Med 160, 521–540. 10.1084/jem.160.2.521.

94. Kaech, S.M., and Ahmed, R. (2001). Memory CD8+ T cell differentiation: initial antigen encounter triggers a developmental program in naive cells. Nat Immunol 2, 415–422. 10.1038/87720.

95. Maekawa, Y., Minato, Y., Ishifune, C., Kurihara, T., Kitamura, A., Kojima, H., Yagita, H., Sakata-Yanagimoto, M., Saito, T., Taniuchi, I., et al. (2008). Notch2 integrates signaling by the transcription factors RBP-J and CREB1 to promote T cell cytotoxicity. Nat Immunol 9, 1140–1147. 10.1038/ni.1649.

96. Leveen, P., Larsson, J., Ehinger, M., Cilio, C.M., Sundler, M., Sjostrand, L.J., Holmdahl, R., and Karlsson, S. (2002). Induced disruption of the transforming growth factor beta type II receptor gene in mice causes a lethal inflammatory disorder that is transplantable. Blood 100, 560–568. 10.1182/blood.v100.2.560.

97. Steinke, F.C., Yu, S., Zhou, X., He, B., Yang, W., Zhou, B., Kawamoto, H., Zhu, J., Tan, K., and Xue, H.H. (2014). TCF-1 and LEF-1 act upstream of Th-POK to promote the CD4(+) T cell fate and interact with Runx3 to silence Cd4 in CD8(+) T cells. Nat Immunol 15, 646–656. 10.1038/ni.2897.

98. Strauss, L., Mahmoud, M.A.A., Weaver, J.D., Tijaro-Ovalle, N.M., Christofides, A., Wang, Q., Pal, R., Yuan, M., Asara, J., Patsoukis, N., and Boussiotis, V.A. (2020). Targeted deletion of PD-1 in myeloid cells induces antitumor immunity. Sci Immunol 5. 10.1126/sciimmunol.aay1863.

99. Stuart, T., Butler, A., Hoffman, P., Hafemeister, C., Papalexi, E., Mauck, W.M., 3rd, Hao, Y., Stoeckius, M., Smibert, P., and Satija, R. (2019). Comprehensive Integration of Single-Cell Data. Cell 177, 1888–1902 e1821. 10.1016/j.cell.2019.05.031.

100. Bunis, D.G., Andrews, J., Fragiadakis, G.K., Burt, T.D., and Sirota, M. (2021). dittoSeq: universal user-friendly single-cell and bulk RNA sequencing visualization toolkit. Bioinformatics 36, 5535–5536. 10.1093/bioinformatics/btaa1011.

101. Picelli, S., Faridani, O.R., Bjorklund, A.K., Winberg, G., Sagasser, S., and Sandberg, R. (2014). Full-length RNA-seq from single cells using Smart-seq2. Nat Protoc 9, 171–181. 10.1038/nprot.2014.006.

102. Hahaut, V., Pavlinic, D., Carbone, W., Schuierer, S., Balmer, P., Quinodoz, M., Renner, M., Roma, G., Cowan, C.S., and Picelli, S. (2022). Fast and highly sensitive full-length single-cell RNA sequencing using FLASH-seq. Nat Biotechnol 40, 1447–1451. 10.1038/s41587-022-01312-3.

103. Dobin, A., Davis, C.A., Schlesinger, F., Drenkow, J., Zaleski, C., Jha, S., Batut, P., Chaisson, M., and Gingeras, T.R. (2013). STAR: ultrafast universal RNA-seq aligner. Bioinformatics 29, 15–21. 10.1093/bioinformatics/bts635.

104. Bray, N.L., Pimentel, H., Melsted, P., and Pachter, L. (2016). Near-optimal probabilistic RNA-seq quantification. Nat Biotechnol 34, 525–527. 10.1038/nbt.3519.

105. Lun, A.T.L., Riesenfeld, S., Andrews, T., Dao, T.P., Gomes, T., participants in the 1st Human Cell Atlas, J., and Marioni, J.C. (2019). EmptyDrops: distinguishing cells from empty droplets in droplet-based single-cell RNA sequencing data. Genome Biol 20, 63. 10.1186/s13059-019-1662-y.

106. Hafemeister, C., and Satija, R. (2019). Normalization and variance stabilization of single-cell RNA-seq data using regularized negative binomial regression. Genome Biol 20, 296. 10.1186/s13059-019-1874-1.

